# Cryo-EM Structure of a Ferrichrome Importer FhuCDB

**DOI:** 10.1101/2021.06.15.448540

**Authors:** Wenxin Hu, Hongjin Zheng

## Abstract

As one of the most elegant biological processes developed in bacteria, the siderophore-mediated iron uptake demands the action of specific ATP-binding cassette (ABC) importers. Although extensive studies have been done on various ABC importers, the molecular basis of these iron-chelated-siderophore importers are still not fully understood. Here, we report the structure of a ferrichrome importer FhuCDB from *Escherichia coli* at 3.4Å resolution determined by cryo electron microscopy. The structure revealed a monomeric membrane subunit of FhuB with a substrate translocation pathway in the middle. In the pathway, there were unique arrangements of residues, especially layers of methionines. Important residues found in the structure were interrogated by mutagenesis and functional studies. Surprisingly, the importer’s ATPase activity was decreased upon FhuD binding, which deviated from the current understanding about bacterial ABC importers. In summary, our studies not only reveal a new structural twist in the type II ABC importer subfamily, but also provide biological insights in the transport of iron-chelated siderophores.

## Introduction

Siderophores are a group of small molecules synthesized and secreted by microorganisms to chelate life-essential iron ions from the environment ^1^. The importance of siderophore molecules are further highlighted by the increasing reports of their “non-classical” functions ^2^. These functions include binding and transporting a variety of metal ions as metallophores ^3,4^, acting as signaling molecules for gene regulation ^5^, protecting bacteria from oxidative stress ^6,7^, as well as providing antibacterial activity as sideromycins ^8,9^. To exercise these functions, siderophores have to be recognized and reimported by specific membrane transporters. For example, in the inner membrane of uropathogenic *Escherichia coli*, there are four such transporter systems known so far: YbtPQ importing yersiniabactin ^10^, FepBDGC importing enterobactin ^11^, FecBCDE importing citrate-based siderophores ^12^, as well as FhuCDB importing hydroxamate-based siderophores ^13^. Although all of siderophore importers belong to the superfamily of ATP-binding cassette (ABC) transporters, the architecture of the four systems are not the same: YbtPQ resembles the fold of type IV exporter ^14^, while the other three seem to be bacterial type II importers.

It is well known that ABC transporters have at least four common domains: two transmembrane domains (TMD) forming a central translocation pathway and two cytosolic nucleotide-binding domains (NBD) providing necessary energy for the substrate translocation via ATP hydrolysis. For type II ABC importers in gram-negative bacteria, periplasmic substrate-binding proteins (SBP) are usually required to specifically bind and deliver substrates to their TMDs. Available high-resolution structures of type II importers, including vitamin B12 importer BtuCDF from *E. coli* ^15–18^, heme importers HmuUV from *Y. pestis* ^19^and BhuUVT from *B. cenocepacia* ^20^, as well as molybdate importer MolBC from *H. influenzae* ^21^, have provided excellent explanation for the general mechanism of substrate import. However, how are different iron-chelated-siderophores imported through related ABC importers is not well understood.

In this study, we functionally characterized the ferrichrome, a hydroxamate-type siderophore molecule, importer FhuCDB complex from *E. coli* and determined its structure at 3.4Å resolution by single particle cryo electron microscopy (cryo-EM). As far as we know, this is the first high-resolution structure of a siderophore importer in the type II ABC importer subfamily. The structure was in the inward-open conformation as the periplasmic side of the transport pathway was closed while the cytoplasmic side was open. The substrate-binding pocket in FhuD (SBP), was fully occupied by loops from FhuB (TMD) as well as several water molecules. In all known type II importer structures, the translocation pathway is formed by homodimers of their membrane subunits. While in FhuCDB, the pathway was in the middle of the FhuB monomer, representing a new type of type II importer structure. Along the surface of the pathway, there were small hydrophobic residues, uncharged polar residues, as well as unique layers of Met residues. With the FhuCDB structure and related functional experiments, our study provides new mechanical insights in the import of iron-chelated-siderophores.

## Results and Discussion

### ATPase activity and transport assay of the Fhu importer in proteoliposomes

To characterize the function of Fhu importer, we overexpressed the full FhuCDB complex as well as the FhuCB subcomplex (only the membrane and cytosolic components), purified them in detergent lauryl maltose neopentyl glycol (LMNG) (**Figure S1**). To perform functional experiments, we incorporated FhuCB into liposomes made of *E. coli* polar extract lipids. The orientation of the reconstituted importers is determined by limited proteolysis of the proteoliposomes using thrombin, followed by western blot using anti-His probe (**Figure S2A**). The idea was that FhuC from the inside-out importers would be digested and lose its N-terminal His-tag, while FhuC from the right-side-out importers would not be digested. The results (**Figure S2B**) showed that ~64% FhuC remained intact after the treatment. Considering that the efficiency of the thrombin digestion was ~90%, thus in the proteoliposomes, ~60% of the reconstituted FhuCB were right-side-out and the rest ~40% of them were inside-out.

These proteoliposomes were used to measure the ATPase activity of the importers. For the FhuCB subcomplex alone, the initial rate of ATP hydrolysis of the inside-out importers in the first 4 min of reaction was measured and appeared to be linear (**Figure S3**). We then plotted the initial rates as a function of the ATP concentration (**Figure 1A**). Here, the ATPase null-mutant (E173A, in the Walker-B domain of FhuC) was used as a negative control as it had no detectable activity. In addition, vanadate was able to inhibit more than 90% of the ATPase activity. While for the wild-type FhuCB, the data were fit using an expanded version of the Michaelis-Menten equation with Hill coefficient, which yielded the following kinetic constants: V_max_ of ~745 ± 31 nmol/mg/min, K_m_ of ~0.69 ± 0.08 mM and Hill coefficient of ~1.97 ± 0.08. When comparing to known Type II ABC importers (**Table S1**), we found that FhuCB had a relatively high K_m_ value, suggesting a lower binding affinity to ATP. Furthermore, the sigmoidal shape of the curve with a Hill coefficient of ~1.97 indicated a strong cooperativity between the two ATP binding sites in FhuC. To test if the importer’s ATPase activity was affected by FhuD, we performed the same experiment using FhuCB proteoliposomes with enclosed ferrichrome-loaded FhuD and FhuD only. Surprisingly, the results (**Figure 1B**) showed that the ATPase activity of the FhuCB + FhuD was decreased as its kinetic constants were V_max_ of ~127 ± 8 nmol/mg/min, K_m_ of ~0.74 ± 0.05 mM and Hill coefficient of ~1.45 ± 0.04. While with the ferrichrome-loaded FhuD, the importer’s ATPase activity was also decreased as the kinetic constants were V_max_ of ~155 ± 19 nmol/mg/min, K_m_ of ~0.77 ± 0.06 mM and Hill coefficient of ~1.73 ± 0.06. The consensus notion in the field is that docking of the SBP to the TMD stimulates the ATPase activity of all known ABC importers ^22^. However, in the case of Fhu importer, upon FhuD binding, its ATPase activity was apparently decreased as the V_max_ dropped more than 5 times. In the meantime, the cooperativity of the ATP binding was weakened, thought the ATP binding affinity seemed to be at the same level as indicated by similar K_m_ values. These results suggest that the molecular mechanism of FhuCDB might deviate from the standard model of type II ABC importers derived from BtuCDF.

**Figure 1.**
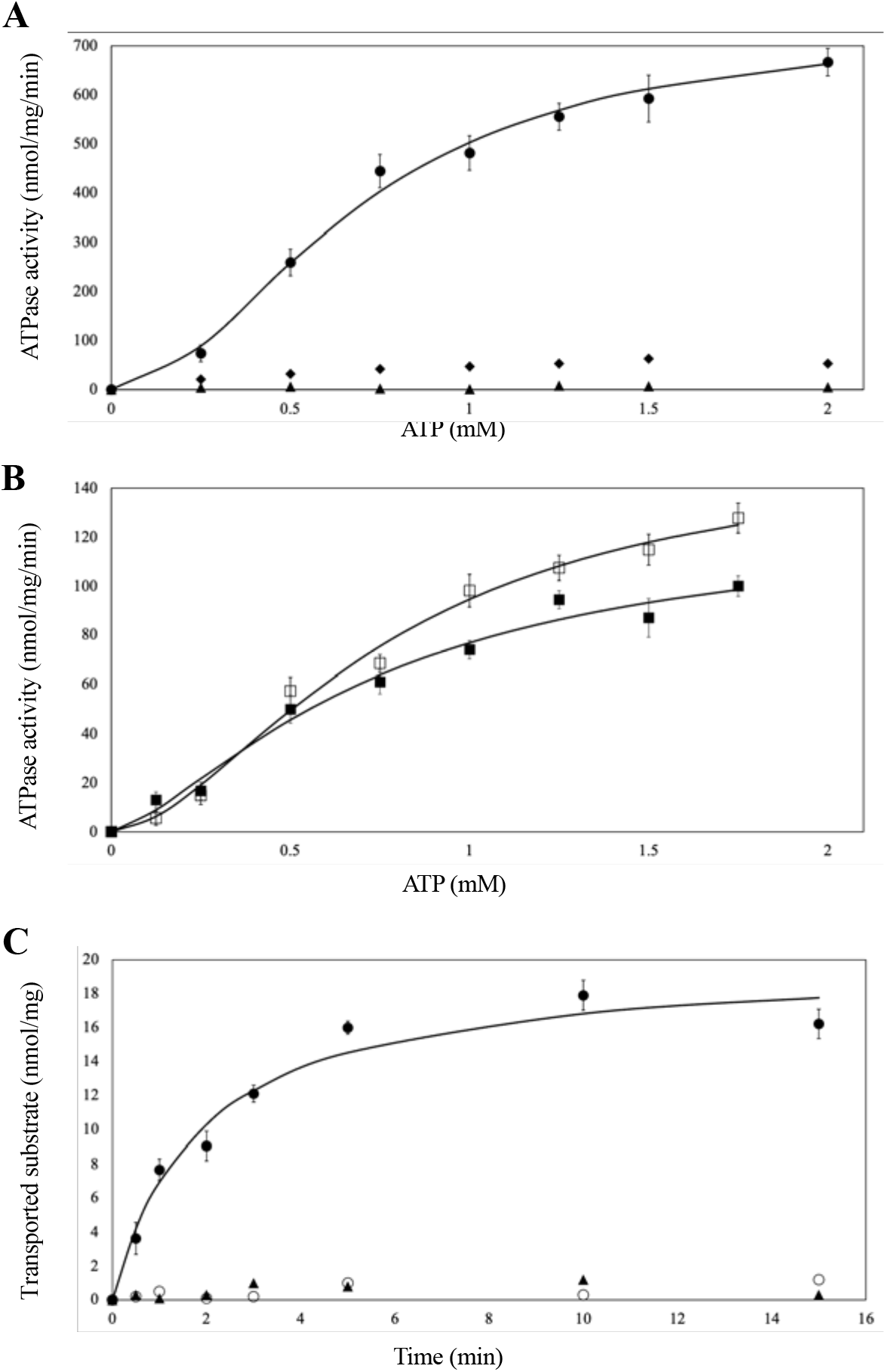
Functional characterization of the Fhu importers in proteoliposomes. **A & B**, ATPase activity of FhuCB plotted against ATP concentrations. **C**, Substrate transport activity of FhuCB in proteoliposomes. The samples are FhuCB (●), vanadate inhibited FhuCB (♦), E173A ATPase null-mutant FhuCB (▲), FhuCB with ferrichrome-loaded FhuD (□), FhuCB with FhuD (■), and FhuCB transport in the absence of FhuD (○). Error bars are the standard deviation from three independent measurements.

For the transport assay, an ATP-regenerating system with ATP and MgCl_2_ was included inside the reconstituted proteoliposomes by following the established protocol ^23^. The transport activity of the right-side-out FhuCB was quantified by liquid scintillation counting as ^55^Fe-ferrichrome was used as the substrate. The results (**Figure 1C**) showed that the ^55^Fe-ferrichrome uptake followed the Michaelis-Menten equation and reached saturation in ~10min. The initial rate of the transport was ~7.5nmol/mg/min. Ideally, the coupling efficiency of the ATPase activity and transportation rate of ABC transporters is 2, meaning two ATP are hydrolyzed when one substrate is translocated. In the case of FhuCDB, if assuming that FhuCDB is the main species *in vivo*, the calculated coupling efficiency is ~20, which is higher than the theoretical value of 2 but reflects the universal characterization of inefficient translocation among most ABC transporters ^24.^

### Overall architecture of FhuCDB in the inward-open conformation

To understand the molecular basis of this importer, we determined the structure of FhuCDB in LMNG by single particle cryo electron microscopy at 3.4Å resolution (**Table S2, Figure S4, S5 & S6**). The overall size of the complex was ~130 x 100 x 80 Å^3^, with one FhuD on the periplasmic side, one FhuB in the membrane and two FhuC on the cytosolic side (**Figure 2A**). The overall fold of the complex indicated that the importer belonged to the type II ABC importer subfamily. However, a unique feature in FhuCDB was the single membrane subunit of FhuB, comparing to two subunits of homodimers in other known subfamily members. FhuB had 20 transmembrane helices (TM) folding into two domains: N-half with the first 10 TMs and C-half with the last 10 TMs (**Figure 2B**). In between the two halves, there was a central translocation pathway. Based on a tunnel calculation using the MoleOnline server ^25^, the FhuB central pathway was open to the cytosolic side and closed to the periplasmic side, indicating an inward-open conformation (**Figure 2A**). Right at the cytosolic opening of the central pathway, there was a loop of ~24 residues (residues 324 ~ 347) connecting TM10 from N-half and TM11 from C-half that are 45Å away from each other. Unfortunately, the cryo-EM density of this loop was mostly missing, suggesting that the loop was fairly flexible. Although the loop was unlikely to block the channel opening in this inward-open conformation, it is unclear at this point whether its local state would change in the occluded or outward-open conformations.

**Figure 2.**
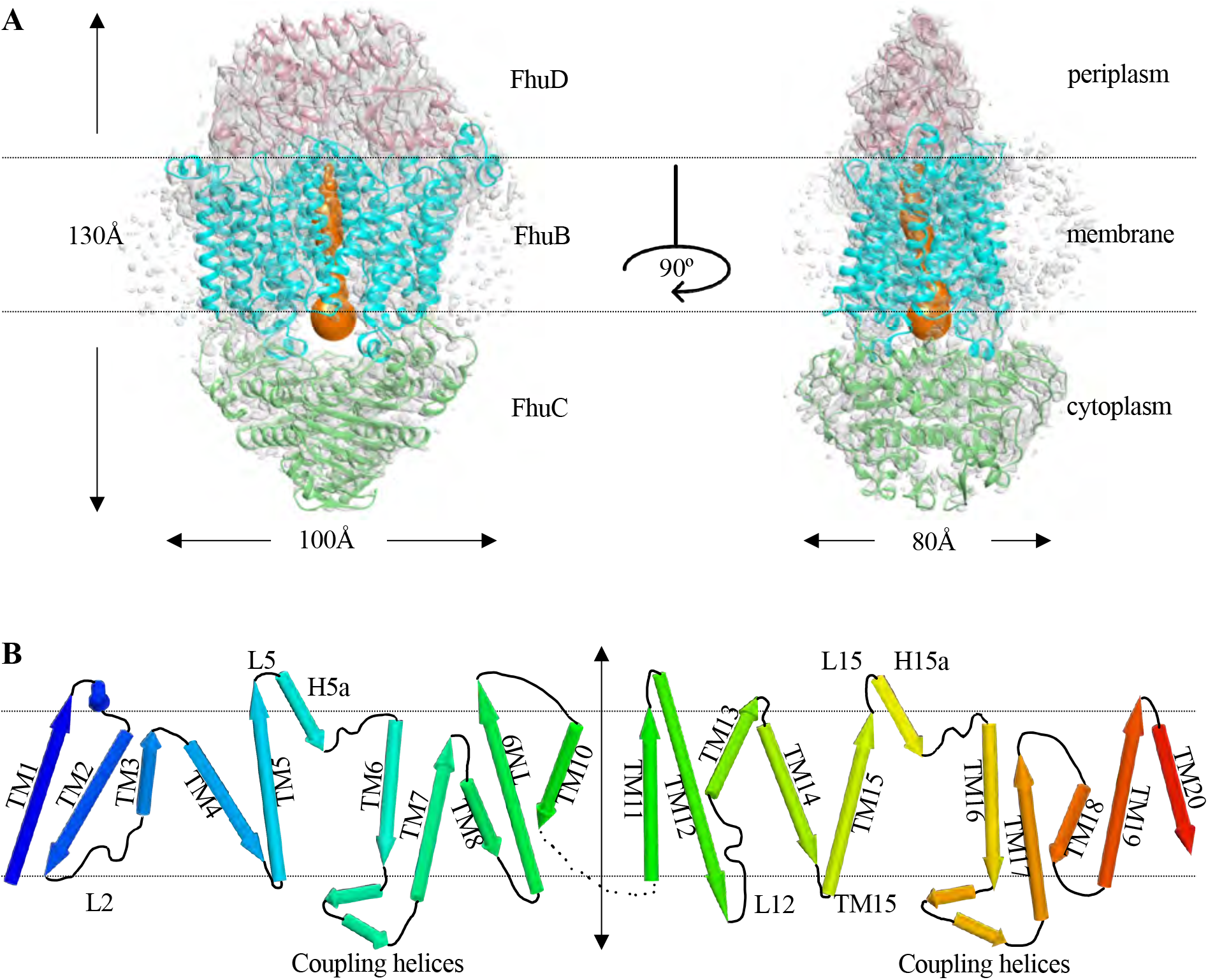
Overall structure of FhuCDB. **A**, Atomic model of FhuCDB fits in the cryo-EM density (gray). FhuD in pink, FhuB in cyan, and two FhuC in green. The calculated tunnel in FhuB (orange) indicates that the complex is in the inward-open conformation. **B**, Topology of FhuB showing 20 TMs colored in rainbow with N-terminal in blue and C-terminal in red. Important helices and loops discussed in this article, as well as all TMs are labeled.

### Substrate-binding site in FhuD occupied by FhuB loops and water molecules

FhuD is a Cluster A type periplasmic SBP for the FhuCDB import system ^26^. It has two lobes (not identical to each other) connected by a hinge α-helix (backbone), which ensures a rigid overall fold and allows only small movement upon substrate binding (**Figure 3A**). In our structure, although there was no substrate bound, FhuD could be superimposed well to all previously determined substrate-loaded FhuD structures ^27,28^ with a root mean square deviation (r.m.s.d) of 2.3~2.5Å (**Figure 3A & S7**). Using the gallichrome-loaded FhuD (PDB:1EFD) as a comparison, the C-lobe of our FhuD moved marginally with most substrate-interacting residues (hydrophobic I183, L189 and W273) remaining in position except W217, which was flipped 180°. In contrast, the N-lobe moved substantially away from the center, leaving only one residue (W68) unchanged while most other substrate-interacting residues (R84, S85, Y106) pulled away. For instance, the OH group of the Y106 side chain moved as far as ~9Å. The movement of these residues effectively deformed the original substrate-binding pocket in FhuD and allowed it to be occupied by two loops (scoop loops) from FhuB: L5 in between TM5 and H5a as well as L15 in between TM15 and H15a (**Figure 3B**). The “scoop loop” was originally described in the crystal structure of maltose transporter MalFGK ^29^, as the P3 loop from MalG clashed into the maltose binding site in MBP. After that, similar loops were found in BtuCDF and BhuUVT complex structures ^17,20^. Thus, it appears to be a common theme for ABC importers to use these loops to scoop out substrates from their SBP and subsequently deliver them to their TMD. In FhuCDB, three residues in L5 (F167, D170, Q171) were directly interacting with FhuD residues (N64, W68) via hydrogen bonding. While L15 loop protruded into the FhuD pocket without apparent interactions. However, an interesting observation here is that there were three structured water molecules. Specially, two of them were trapped in the interface by the hydrophobic bubble created by L15 and the surrounding FhuD loops. Although such water molecules had been observed in multiple MBP-MalFGK crystal structures, they had never been described in type II importers such as BtuCDF. It is possible that the water molecules are critical for the association of FhuD and FhuB ^30^.

**Figure 3.**
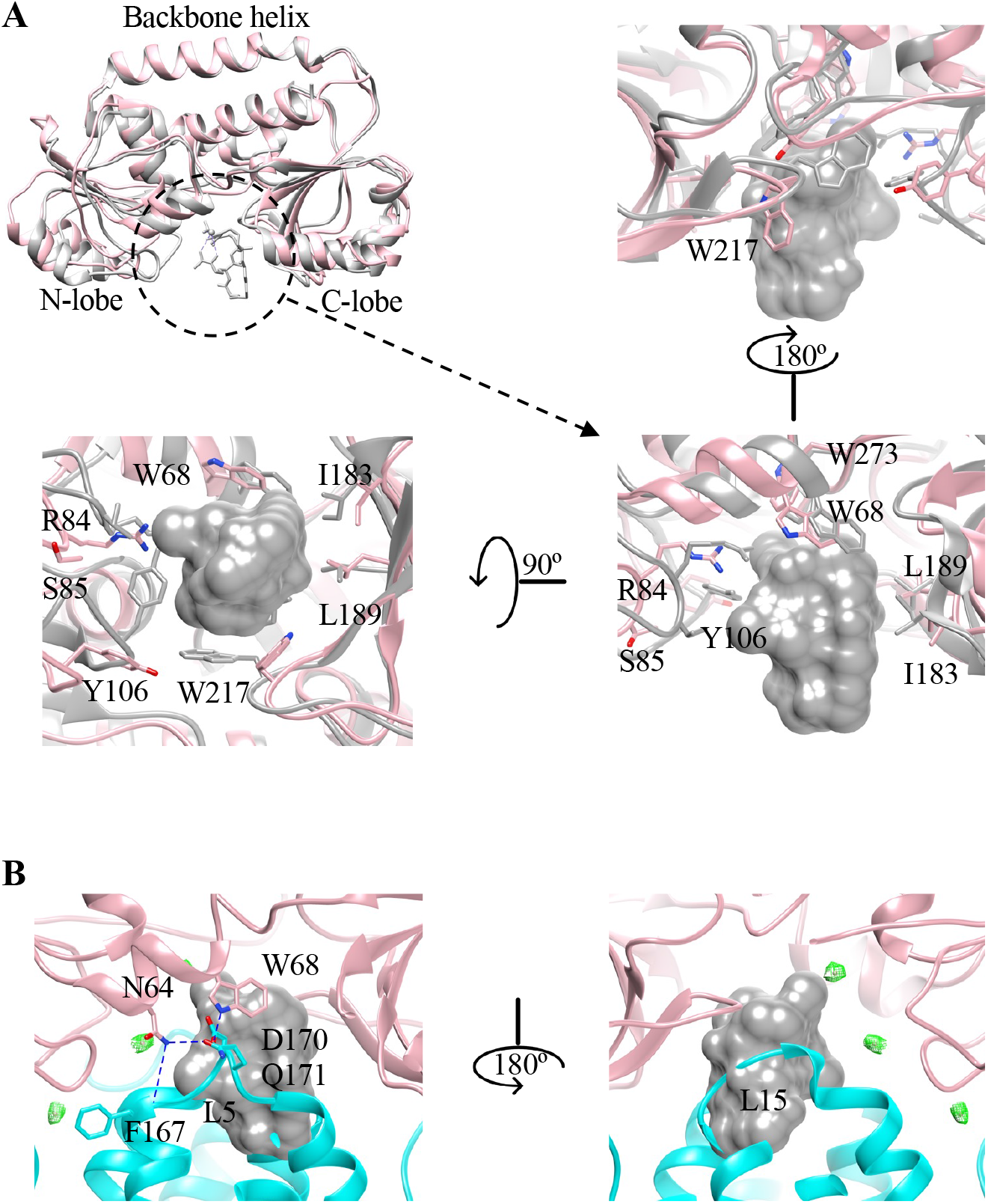
Deformed substrate-binding pocket in FhuD. **A**, Superimposed FhuD structures from FhuCDB (pink) and gallichrome-bound FhuD (PDB:1EFD, gray) with substrate-binding residues shown in zoomed-in views. **B**, Substrate-binding pocket in FhuD is occupied by two FhuB loops L5 and L15 (cyan) with specific hydrogen bonds (blue). Experimental densities of trapped water molecules are shown in mesh (green).

### Subunit interactions within FhuCDB

The rearranged FhuD substrate-binding pocket was not the only communication between FhuD and FhuB in the structure. In fact, FhuD docked onto FhuB through a complicated network of interactions. On one hand, the N-lobe of FhuD docked onto the C-half of FhuB via the following specific interactions FhuD-E90/FhuB-R390, FhuD-N88/FhuB-Q636, FhuD-N88/FhuB-Y517, FhuD-E86/FhuB-S515, as well as a hydrogen bonding network involving a third water molecule (**Figure S8A**). This water not only connected FhuB-T510 and the backbone of FhuD-T85, but also stabilized the side chain of FhuB-Q507 which in turn interacted with the backbone of FhuB-G107. On the other hand, the C-lobe of FhuD docked onto the N-half of FhuB involving the following FhuD residues H187, S223, D225, R226 and FhuB residues S57, T182, T184, E304 (**Figure S8B**). Most FhuB residues interacting directly with FhuD were highly conserved (**Figure S9**). When we divided the whole FhuCDB complex into two parts along the central pseudo two-fold axis, we found that the two FhuD lobes were completely misaligned, the two FhuB halves were superimposed reasonably well, while the two FhuC subunits were almost identical (**Figure S10A**). The association between FhuB and FhuC was symmetrical: the coupling helices from FhuB-N-half (residues 209~226) matched well with the coupling helices from FhuB-C-half (residues 542~559). Similar to other ABC transporters, the coupling helices docked into the surface grooves of ATPase through mainly hydrophobic interactions and a few hydrogen bonds. It was reported that mutating two conserved Gly residues (G226 and G559) to Ala, Val or Glu reduced iron hydroxamate uptake ^31^. Our structure explained it well, as there was no room for any side chains at this position, thus these mutations would push away the adjacent helix on FhuC and destroy the strong association between the two proteins (**Figure S10B)**.

To further understand the association between subunits, we measured the binding affinity between FhuCB and FhuD in detergent by microscale thermophoresis (MST) ^32^. The results (**Figure S11**, **Table S3**) showed that, without substrate, the binding between FhuCB and FhuD was reasonably tight as the K_d_ values were in the μM range no matter ATP-Mg is present or not. Considering that FhuCDB were overexpressed in rich media, the amount of synthesized siderophore molecules in *E. coli* was negligible. Thus, without *in vivo* substrates, the naturally strong interaction between FhuCB and FhuD was most likely the reason why we were able to purify FhuCDB as an intact complex. When FhuD was loaded with ferrichrome, the binding was dramatically increased, as the K_d_ values dropped to ~12.2 nM (without ATP-Mg) and ~1.03 nM (with ATP). In addition, we mutated all previously mentioned residues in FhuB (R390, Q636, Q507T510, S515Y517, D170Q171, S57, E304, T182T184) to Ala and found decreased binding affinities between FhuB mutants and FhuD-ferrichrome all across the board (**Table S3**). These measurements agreed with a previous report that both FhuD alone and substrate-loaded FhuD were able to interact with FhuB while tested by chemical cross-linking and by prevention of FhuB degradation by added FhuD ^33^.

### Potential translocation pathway and the gating

In all known type II importers, two TMs along the translocation pathway have been extensively discussed: TM5 on one protomer and its counterpart TM5’ on the other protomer. Comparing to the structure of inward-open BhuUVT (PDB:5B58), the translocation pathway in FhuB was vividly narrower as TM15 moves ~4.5Å towards the center while other adjacent TMs stayed at similar positions as in BhuU (**Figure S12**). The pathway in FhuB was thus unsymmetrically formed by three main TMs: TM3, TM5 and TM15, without TM13 as in BhuUVT (**Figure 4**). On the periplasmic side, the pathway was closed by residues from the following regions: TM5~H5a and TM15~H15a. Specifically, these four residues: V165, F176 on the N-half and their counterparts Q498, L509 on the C-half, defined the narrowest position in the pathway, which was then directly closed on top by H169 and M505 (**Figure 4A**). Below the periplasmic gate, residues along the path could be grouped into three types: hydrophobic residues with mostly small side chains (A96, G150, L151, L155, G158, A159, L494, I512), polar residues (T91, T92, T97, Q100, S154, N161, Q162, S179, S421, E423, S487, T488, T491, S513) capable of forming hydrogen bonds, as well as Met residues (M175, M495, M492, M484) (**Figure 4B**). How do the specific arrangements of residues contribute to the translocation of substrate such as ferrichrome? First, it has been demonstrated that ferrichrome favorably interacts with aromatic rings and thus there are adequate aromatic residues found in the ferrichrome-binding site in FhuD and its outer membrane receptor FhuA ^30,34^. While in FhuB, no aromatic residues are found pointing directly towards the central pathway, suggesting that the binding between FhuB and ferrichrome will be rather weak and thus favors rapid translocation. Second, to minimize the energy barrier of transportation, oligosaccharide transporters, such as the maltoporin channel ^35^ and Wzm-Wzt transporter ^36^, employ a combination of aromatic hydrophobic interactions with a continuous pattern of hydrogen bond donors and acceptors along their translocation pathways. While in the case of FhuCDB, ferrichrome is not inherently hydrophobic, especially its hexa-peptide ring. Thus, its translocation should be facilitated by the continuous layers of hydrogen bond doners and acceptors along the way, which are the abundant polar residues especially Thr and Ser. Third, it has been shown that chelated Fe^3+^, for instance heme, favors to interact with axially positioned His and Met ^37,38^. In FhuB, the sequentially positioned Met residues along the pathway may transiently interact with Fe^3+^ in the ferrichrome to drive down the substrate. Interestingly, a similar arrangement of Met residues is also observed in another siderophore ABC importer YbtPQ ^14^ (**Figure S13**). To confirm the functional importance of the Met residues along the pathway, we mutated all four of them (M175, M484, M492, M495) to Ala and performed ^55^Fe-ferrichrome transport assay using proteoliposomes. The results show that the Met mutant FhuCB reconstituted into the liposomes as efficiently as wild-type FhuCB, while the transport efficiency drops ~80% (**Figure S14**).

**Figure 4.**
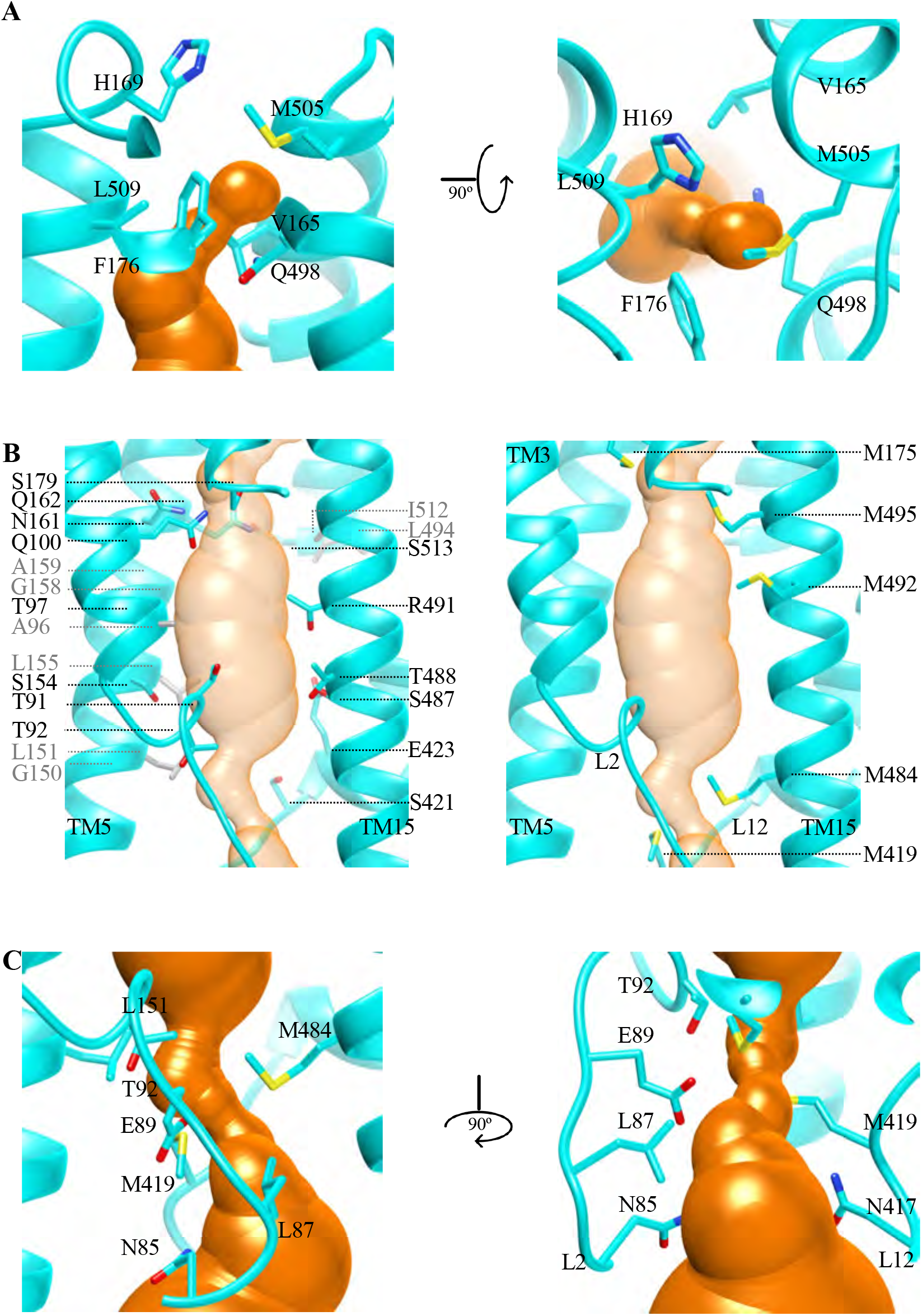
The translocation pathway and its gating in FhuB. **A**, The closed periplasmic gate of the translocation pathway (orange) defined by specific residues (cyan). **B**, Three groups of residues along the translocation pathway from three main TMs: 3, 5 and 15. The left panel shows two groups: small hydrophobic residues (gray) and polar residues (cyan), while the right panel shows continuous layers of Met residues (cyan). **C**, The narrowest position in the cytoplasmic side of the pathway defined by specific residues.

Previous studies on BtuCDF complex revealed two cytoplasmic gates: gate I of the TM5 that was closed in the structure of asymmetrically closed BtuCDF (PDB: 2QI9) ^16^, and gate II of the L2 loop between TM2 and TM3 that was closed in the structure of outward-open BtuCD (PDB: 4R9U) ^18^. Here, we compared those two structures with our FhuCDB (**Figure S15**). The comparison showed roughly the same position for the cytosolic half of these TMs: FhuB-TM15 and BtuC-TM5’. However, the positions of TM3, TM5 and L2 in FhuB were drastically different. Specifically, FhuB-TM3 was much closer to the center of the molecule than BtuC-TM3 in both Btu structures, and thus actively participated in the formation of the translocation pathway. As cytoplasmic gate I, the cytoplasmic half of FhuB-TM5 was tilted away from the center, making it different from the closed TM5 in the 2QI9 structure (**Figure S15A**) but similar to the open TM5 in the 4R9U structure (**Figure S15B**). Thus, the cytoplasmic gate I was open in our structure. In the meantime, the cytoplasmic gate II in FhuB was also open, because the distance between the L2 and L12 loops were much larger than the closed gate II in the outward-open BtuCD (**Figure S15B**). The closed gate II in BtuCD was formed by two conserved residues N83 and L85, equivalent to residues N85, N417, and L87, M419 in FhuB, respectively. While in our structure, the narrowest position, which is still open, in the cytoplasmic side of the pathway was defined by residues T92, M484, L151 and E89 (**Figure 4C**).

## Conclusion

In this study, we determined the structure of the ferrichrome ABC importer FhuCDB from *E. coli* in the inward-open conformation at 3.4Å resolution. The structure revealed critical residues regarding subunit interactions, periplasmic gating, as well as transport pathway. The importance of these residues was confirmed by mutagenesis and functional assays. Specifically, along the transport pathway, a continuous layer of Met residues along the transport pathway seemed to be a unique feature of the siderophore ABC importers.

By comparing with other known type II importers, we propose the following molecular mechanism for the substrate import. First, the FhuCDB structure we obtained most likely represent the resting state between substrate import cycles because of the following three reasons: 1) there is no natural substrate synthesized in *E.coli* growing in rich medium with high iron concentration, and thus no import event happened; 2) FhuD stays on FhuCB despite the natural high ATP concentration in the cells; 3) FhuCDB is on the same operon with overlapping nucleotides between adjacent genes, thus the co-translation of the three proteins should promote the formation of the intact complex. Furthermore, we incubated the purified FhuCDB complex at 1μM with 10μM ferrichrome and 5mM ATP-Mg in the solution followed by gel filtration analysis and found no free FhuD, suggesting that neither ATP nor substrate was enough to release FhuD from FhuCB. Thus, the only way to start the substrate import seems to be competitively replacing FhuD with substrate-loaded FhuD. This is possible because substrate-loaded-FhuD has higher binding affinity to FhuCB as shown in our experiments. However, for vitamin B12 importer BtuCDF: without substrate, BtuF binds to BtuCD extremely strongly (K_d_ = 0.12pM), while substrate-loaded BtuF still binds to BtuCD but much weaker (K_d_ = 21.1nM without ATP-Mg and K_d_ = 3.6nM with ATP-Mg) ^39^. It is clearly an opposite trend of the SBP-TMD association comparing to our study. Next, when FhuD is replaced by substrate-loaded FhuD, with the action of scoop loops, FhuD opens the pocket and release the substrate into the central translocation pathway in FhuB. Here, ATP binding in FhuC should happen in order to change the FhuB conformation to open its periplasmic gate and close its cytoplasmic gates. Then, the substrate is translocated down the pathway via transient interactions with layers of Met and small polar residues. At the same time, ATP hydrolysis triggers FhuB to open its cytoplasmic gates and release the substrate into the cytoplasm. Last, as illustrated in the model derived from BtuCDF, a general idea is that SBP comes off from TMD to prepare the importer for the next cycle of action. In the case of FhuCDB, we argue that FhuCB might be shortlived or even not existed *in vivo*, and thus FhuD shall not be released in between the import cycles. There are at least three reasons for this argument: 1) the ATPase activity of importer is much lower than with FhuD bound, suggesting that FhuCDB is conformationally more rigid; 2) FhuD could not be separated from FhuCB by incubating with ATP and substrates; 3) in our cryo-EM studies on FhuCDB, we did not find any 2D averages of FhuCB alone (without FhuD), indicating low possibility of its physiological existence. Nevertheless, structures of Fhu importers in other conformations and in lipid-surrounded environment determined in the future will provide more insights into its specific molecular mechanism.

## Materials and Methods

### Plasmids

For the FhuCDB construct, the whole operon of FhuCDB was cloned from *Escherichia coli* (*E. coli*) genome into a pET15b (Novagen) vector with a His-tag and a thrombin digestion site on the N-termini of FhuC. For the wild-type FhuCB construct, FhuD gene in the middle of the operon was deleted directly from the FhuCDB-pET15b construct. All mutants of FhuCB used in this manuscript were modified on this FhuCB-pET15b construct by direct mutagenesis. For the FhuD construct, the gene was amplified and cloned into a pET15b vector.

### Protein expression and Purification

FhuCDB and FhuCB constructs were over-expressed in *E. coli* strain of BL21(DE3) C43 (Sigma-Aldrich) at 18°C overnight with 0.2mM Isopropyl β-D-1-thiogalactopyranoside (IPTG, UBPBio). The culture was harvested and resuspended in Buffer A (20mM HEPES at pH 7.5 and 150mM NaCl) with 1mM phenylmethylsulfonyl fluoride (PMSF). The cells were lysed by passing through the M110P microfluidizer (Microfluidics) two times at 15,000psi. Cell debris was removed by centrifugation at 15,000g for 30min. Then, the membrane fractions were collected by ultracentrifugation at 150,000g for 2hrs, resuspended in Buffer A and stored at −20°C. To solubilize the complexes in the membrane, 1% lauryl maltose neopentyl glycol (LMNG) was added and the mixture was incubated at 4°C for 2hrs before ultracentrifugation at 150,000g for 1hr. Proteins in the supernatant were purified by the affinity chromatography with TALON (Clontech) resin, followed by gel filtration chromatography on a Superose 6 column (GE Healthcare Life Sciences) in Buffer A with 0.001% LMNG. The eluted peak fractions were combined and concentrated to ~5mg/ml for further experiments, respectively. FhuD was expressed and purified by following a published protocol ^28^.

### Reconstitution of FhuCB and FhuCDB into proteoliposomes

The experiment was done by following a previously published protocol ^23^. Briefly, 10mg *E. coli* polar extract lipids (Avanti Polar Lipids) were dissolved in chloroform, dried by nitrogen gas, and immediately resuspended in Buffer A to a final concentration of 10mg/ml. The large unilamellar liposome vesicles were made by extruding the suspension through a 400nm polycarbonate membrane filter using a mini extruder (Avanti Polar Lipids). To reconstitute proteoliposomes, the liposomes were destabilized by 0.015% Triton X-100 and then incubated with purified proteins (FhuCDB or FhuCB or FhuCB) at a ratio of 100:1 (wt/wt) for 20min at room temperature. A total of 100mg Bio-Beads SM-2 (Bio-Rad Laboratories) was then added to the mixture to absorb all detergents overnight at 4°C. The reconstituted proteoliposomes were harvested by ultracentrifugation at 180,000g for 20min for further experiments.

### Thrombin digestion of the proteoliposomes

To estimate the ratio of inside-out and right-side-out transporters in the proteoliposomes, thrombin digestion was performed. The FhuCDB and FhuCB proteoliposomes were incubated with human α-thrombin (Enzyme Research Laboratories) at a molar ratio of 100:1 at room temperature for 2hrs, and then subjected to western blot analysis using HisProbe-HRP conjugate (Thermo Fisher Scientific). The density of the protein bands was quantified using GelAnalyzer 19.1 (www.gelanalyzer.com).

### ATPase activity assay

The ATPase activity was determined using an ATPase/GTPase Activity Assay Kit (Sigma-Aldrich) featuring a detectable fluorescent product from malachite green reacting with released phosphate group. Three proteoliposome samples were used: FhuCB, FhuCB with 10μM ferrichrome and 1μM FhuD enclosed, as well as FhuCB with 1μM FhuD enclosed. 0.002~0.005mg/ml of these proteoliposomes were incubated with ATP-MgCl_2_ at various concentrations from 0 to 2mM at 37°C. Several time points between 0 and 4min were taken. The amount of phosphate produced by inside-out transporters was detected by absorbance at 620nm. The data were fitted to the extended Michaelis-Menten equation with Hill coefficient in Excel (Microsoft).

### Transport assay

Proteoliposomes with either FhuCB or FhuCB Met mutant were used, and the transport rate of the right-side-out importers were measured here. Briefly, to enclose the ATP-regeneration system, the proteoliposomes, were subjected to three cycles of freeze and thaw with 10mM ATP, 2mM MgCl_2_, 0.1mg/ml pyruvate kinase and 10mM phosphoenolpyruvate followed by 5 times of passing through a 400nm polycarbonate membrane filter. The proteoliposomes were harvested by ultracentrifugation at 180,000g for 20min, and then resuspended into Buffer A to a final protein concentration of 0.002mg/ml. The uptake experiments were carried out at 37°C with 1.6μM purified FhuD and 16μM substrate ^55^Fe-ferrichrome. The reaction was stopped at different time points by adding 200μl of ice-cold Buffer A, followed by filtration using pre-wetted cellulose nitrate filters. The filters were washed with 1 ml of Buffer A, dried for 1hr, and then dissolved in 3ml of Filter Count scintillation liquid (Perkin Elmer). The ^55^Fe radioactivity within the proteoliposomes was quantified by using a Tri-carb 2910 TR Scintillation counter (Perkin Elmer).

### Microscale thermophoresis

To measure the binding affinity between FhuD and FhuCB, MST experiments were performed with a Monolith NT.115pico (NanoTemper). A standard protocol was followed ^32^. Briefly, the His-tag on purified FhuCB in detergent was labeled with the red-tris-NTA 2^nd^ generation Dye. For each reaction, 40nM FhuCB was mixed with a serial dilution of 1.6μM FhuD with substrate ferrichrome or 30μM FhuD without substrate. Micro thermophoresis was performed using 20% LED power and medium MST power. K_d_ values were all calculated using the NanoTemper software with default settings.

### Cryo-EM structure determination and model building

3μl of purified FhuCDB in LMNG at ~1.2mg/ml were applied to a plasma-cleaned C-flat holy carbon grids (1.2/1.3, 400 mesh, Electron Microscopy Services). The grid was prepared using a Vitrobot Mark IV (Thermo Fisher Scientific) with the environmental chamber set at 100% humidity and 4°C. The grid was blotted for 3.5s and then flash frozen in liquid ethane cooled by liquid nitrogen. Cryo-EM data were collected in the Pacific Northwest Cryo-EM Center (PNCC) on a Titan Krios (Thermo Fisher Scientific) operated at 300keV and equipped with a K3 direct detector (Gatan) together with a Bioquantum energy filter (slit width of 20eV used). A total of 9142 movies were recorded with a pixel size of 0.399Å under super resolution mode, a defocus range of −1μm to −2.5μm, and a total dose of ~65 electrons/Å^2^ over 60 frames. The total exposure time was ~2.1s with a dose rate of ~19.6 electrons/pixel/s. The data were processed using both cryoSPARC v2.15 ^40^ and Relion 3.1 ^41^. Specifically, in cryoSPARC, movies were processed with patch motion correction and patch CTF estimation. ~2000 particles were manually picked, and 10 class averages were generated by 2D classification. 5 good class averages were selected as templates for auto picking. Following standard procedure of particle extraction, 2D classification, ab-initio reconstruction and heterogeneous refinement, best groups of particles were selected for a final reconstruction at 3.4Å resolution. The reconstruction and selected particles were then imported into Relion for further 3D classification and 3D auto-refine. The final reconstruction from Relion was at the same resolution as the one from cryoSPARC. In both programs, the gold-standard FSC curves with a 0.143 cutoff were used to determine the resolution. The model building process was carried out in Coot ^42^ using the map from Relion. X-ray structure 1ESZ ^27^ was used as the initial model to build FhuD, while other part of the FhuCDB model was built manually from the scratch. Briefly, secondary structures were built with the guild from multiple secondary structure prediction programs: Jpred ^43^ and PSSpred ^44^, as well as the well-known topology of the NBDs. Flexible loops were manually built after placing the secondary structures. The final model was refined in PHENIX ^45^. The quality of the model was assessed by MolProbity ^46^. Statistical details could be found in Table S2. All figures were prepared using the program UCSF Chimera ^47^.

## Acknowledgements

We thank the staff in the Pacific Northwest Center, especially Mr. Theo Humphreys, for Cryo-EM data collection (supported by NIH Grant U24GM129547).

## Author contributions

Hu, W. and Zheng, H. designed the experiments, collected and analyzed the data, and wrote the manuscript.

## Data and material availability

Cryo-EM map of FhuCDB was deposited in the Electron Microscopy Data Bank under the accession code EMD-23251. Coordinates of the atomic model of FhuCDB was deposited in the Protein Data Bank under the accession code 7LB8. All other data are available from the corresponding author upon reasonable request.

---

**Figure S1.**
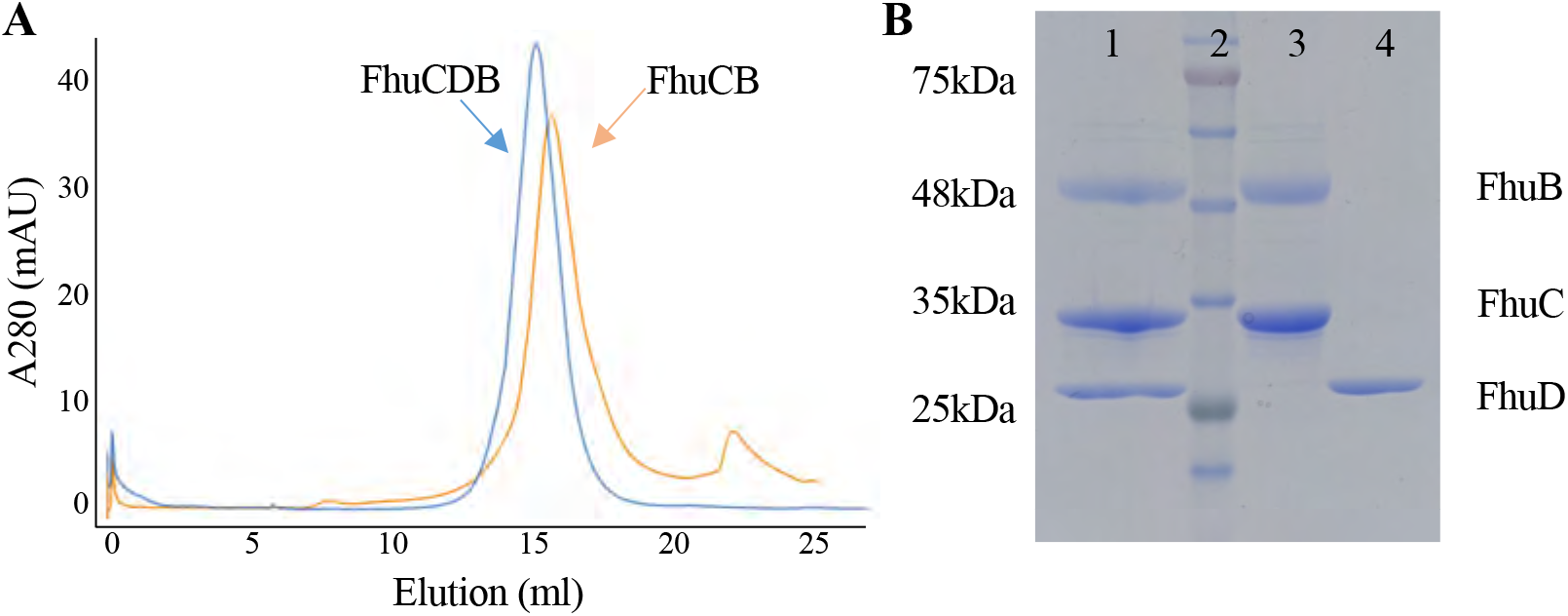
Purification of the Fhu importer. **A,** Gel filtration profiles of the purified FhuCB (orange) and FhuCDB (blue) in detergent LMNG on a S6 column. **B**, SDS-PAGE of the purified FhuCDB (lane 1), FhuCB (lane 3) and FhuD (lane 4). The molecular weight ladder is shown in lane 2.

**Figure S2.**
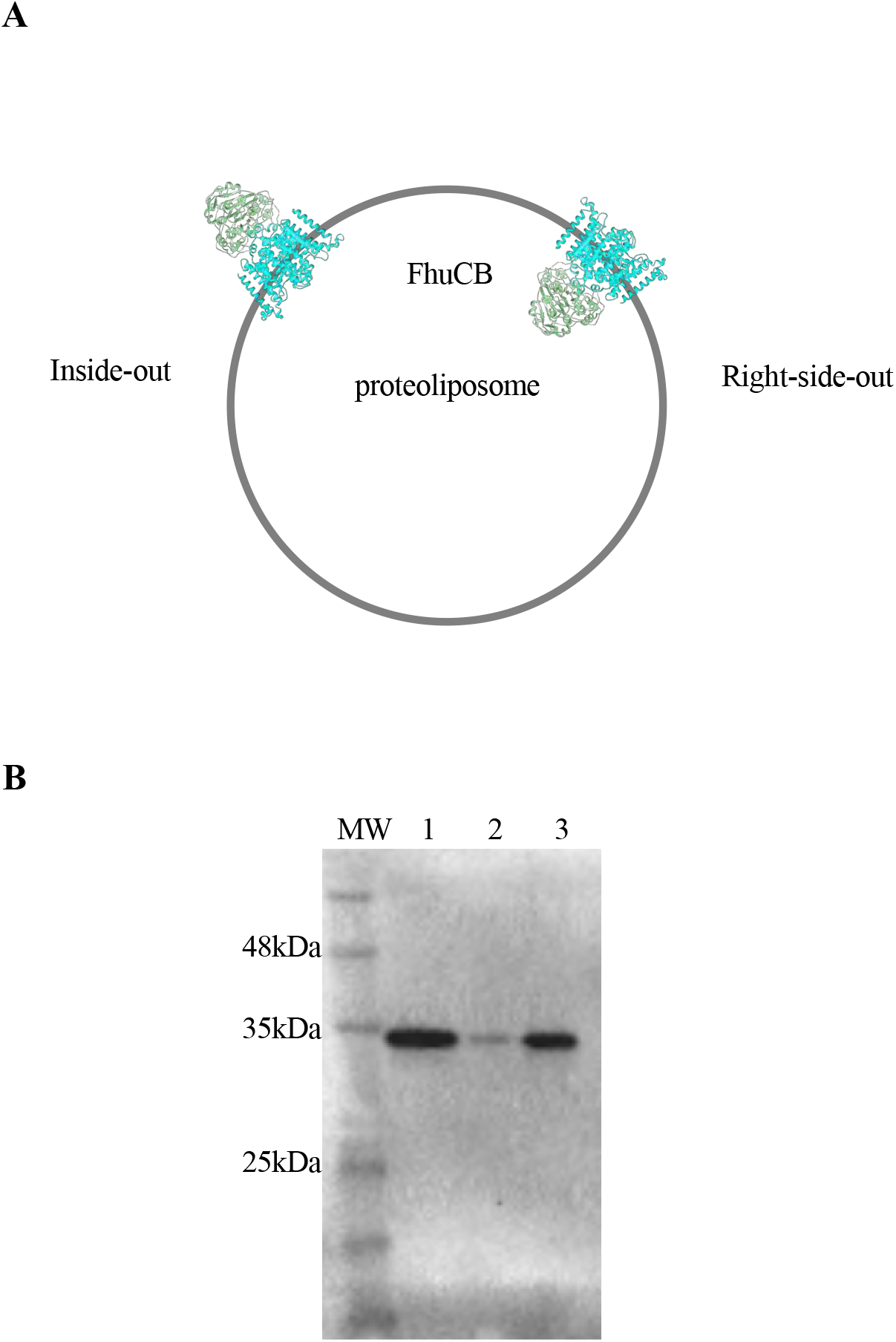
Orientational analysis of the reconstituted FhuCB. **A,** A cartoon representation of a proteoliposome (gray) with two possible orientations of the importers. **B**, Western blot of the thrombin-treated proteoliposomes. Lane 1: untreated FhuCB; lane 2: treated FhuCB in detergent: lane 3: treated FhuCB.

**Figure S3.**
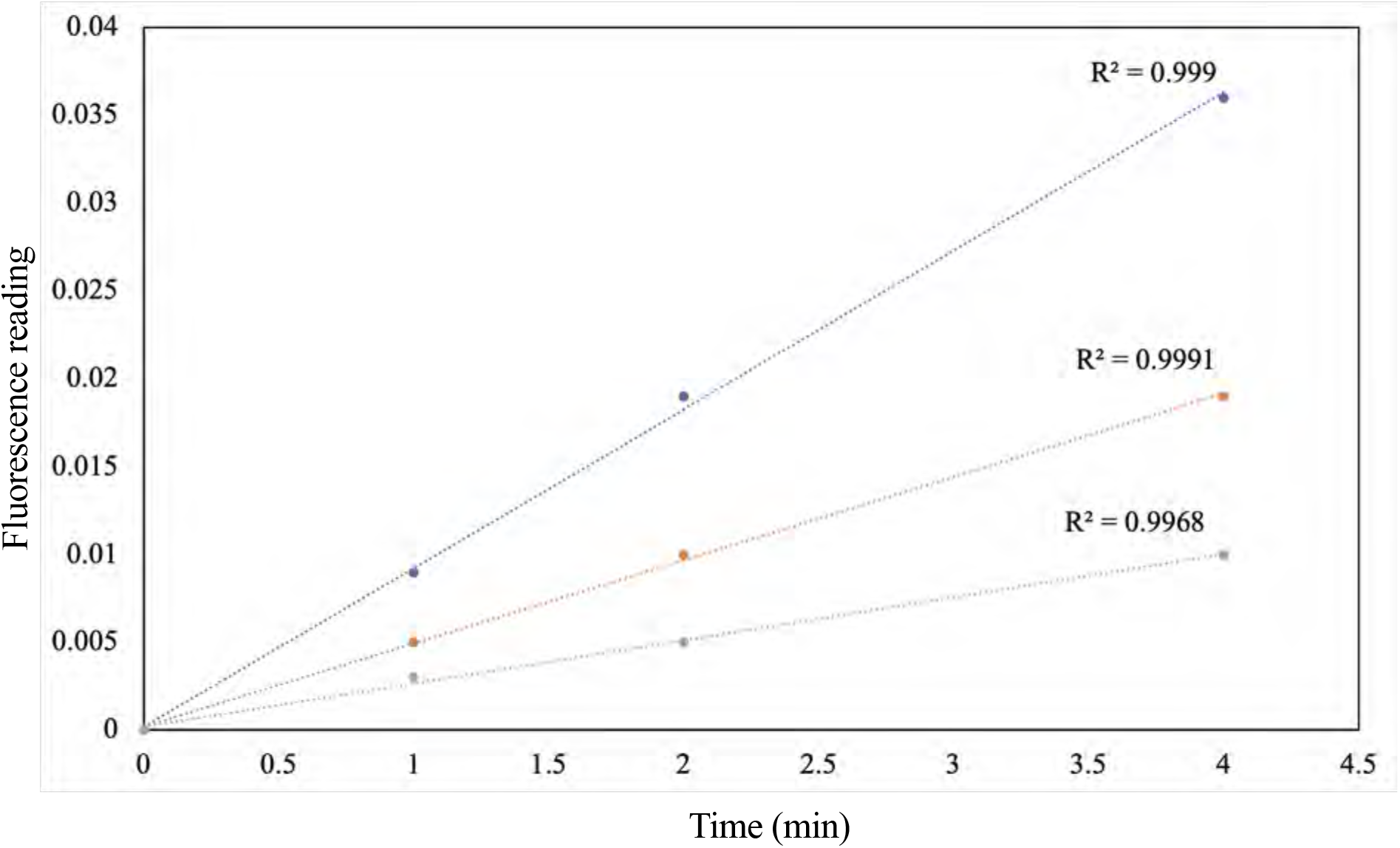
ATPase activity of the FhuCB in proteoliposomes. The initial ATP hydrolysis rates of FhuCB over the first 4 min of reaction at different ATP concentrations of 2mM (blue), 1mM (orange) and 0.5mM (gray). The calculated R^2^ values of each linear fit are shown.

**Figure S4.**
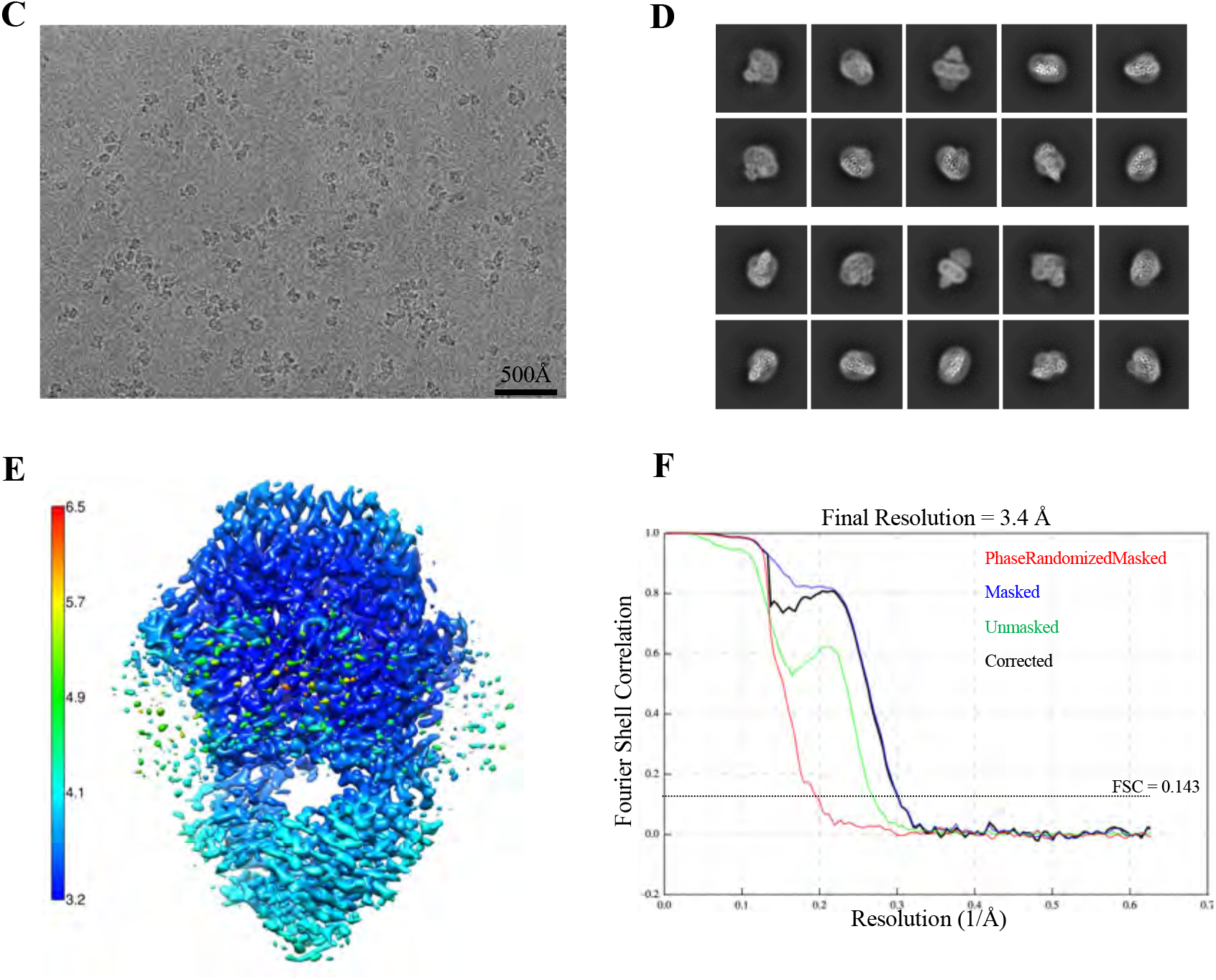
cryo-EM analysis of FhuCDB. **A**, A representative cryo-EM image of FhuCDB. **B**, 2D class averages of FhuCDB. **C**, the final reconstruction of FhuCDB colored by local resolution estimation calculated by Relion. **D**, 3.4Å resolution of the FhuCDB final reconstruction indicated by the gold-standard FSC curve.

**Figure S5.**
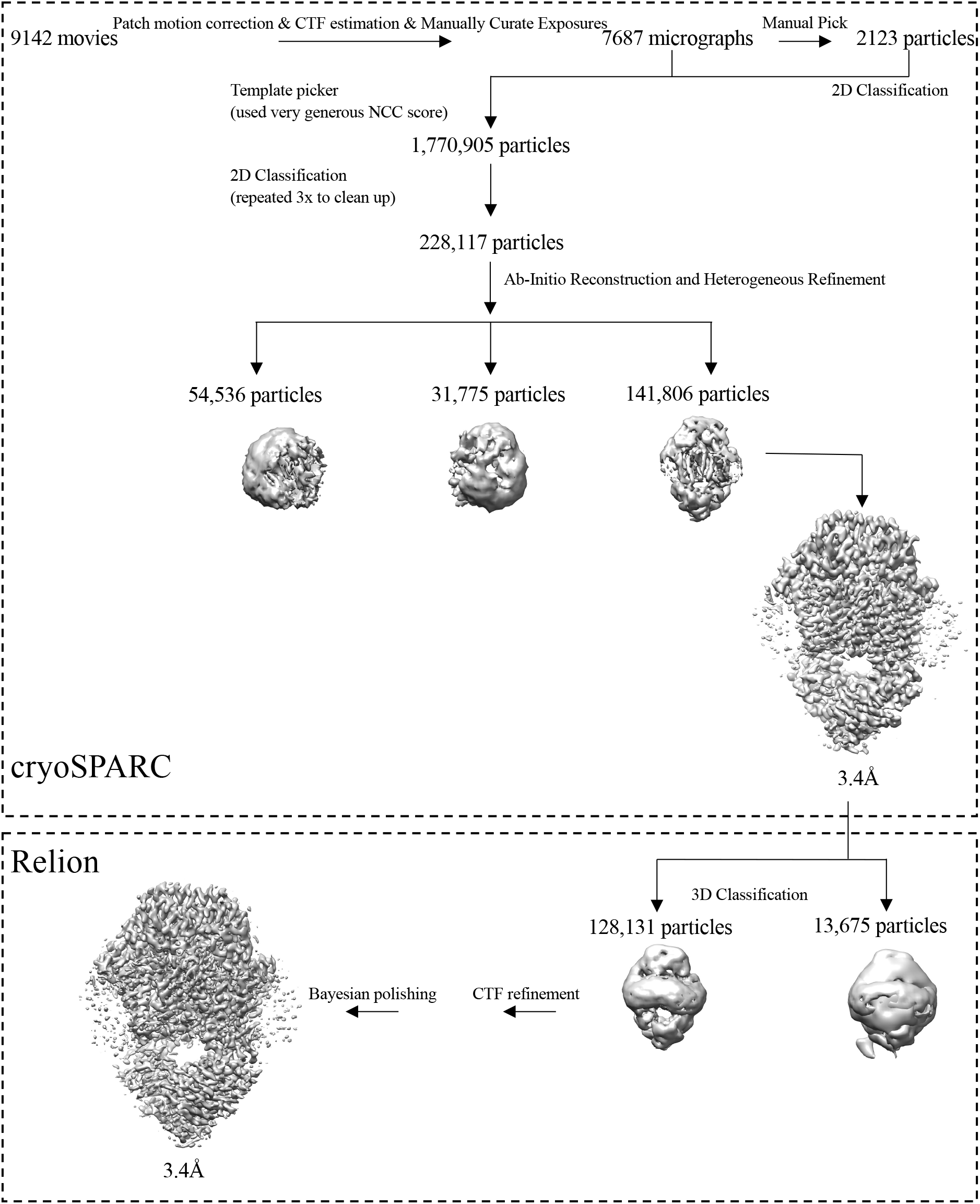
Cryo-EM data processing workflow. The data were processed in both cryoSPARC and Relion software packages. Final reconstructions from both software are at the same 3.4Å resolution.

**Figure S6.**
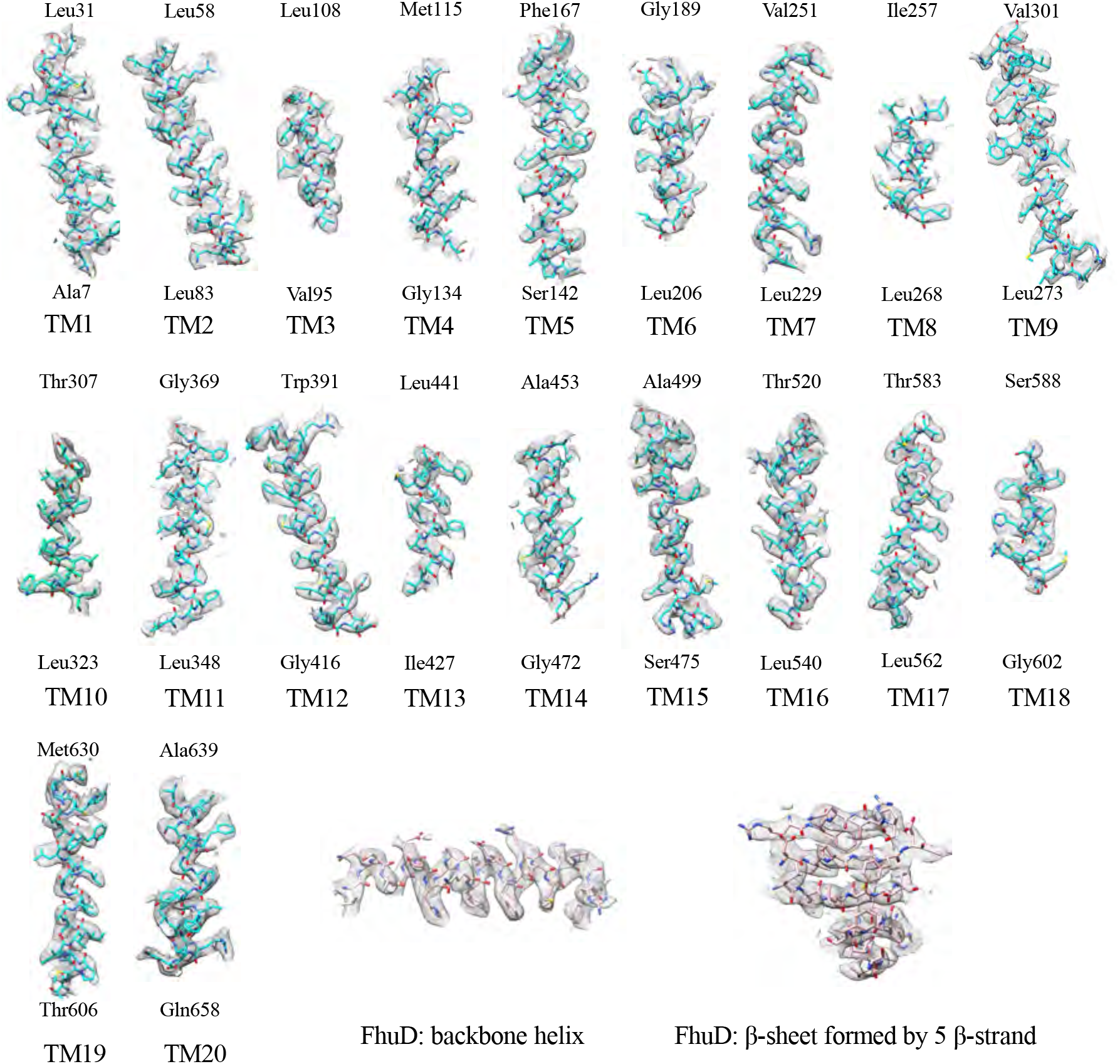
FhuCDB model fits in the experimental density. To demonstrate the quality of the cryo-EM map, part of the atomic model of FhuCDB is overlayed with the density. These include all 20 TMs from FhuB (cyan sticks, with starting and ending residues labeled), as well as the backbone helix and the β-sheet with 5 β-strand of FhuD (pink sticks).

**Figure S7.**
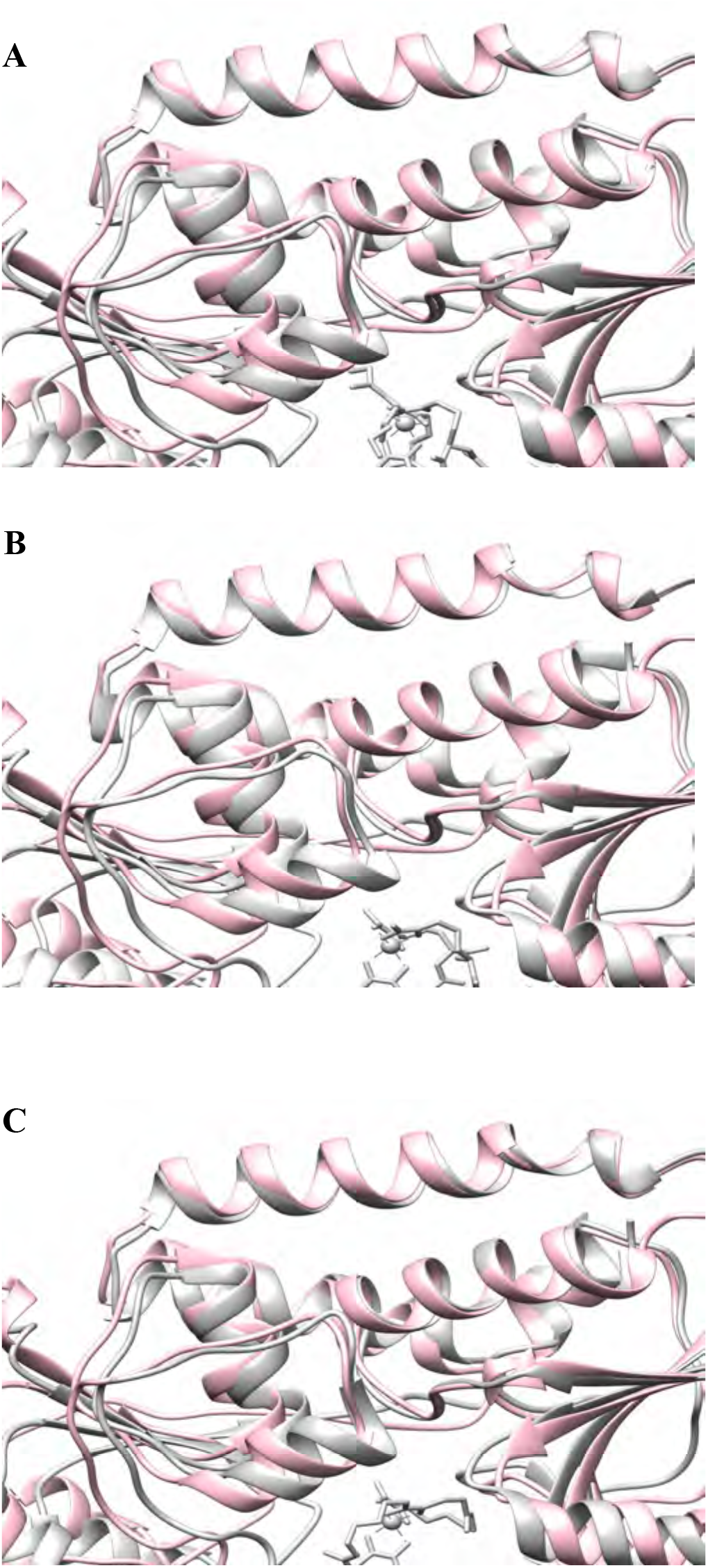
Superposition of FhuD structures. FhuD in the FhuCDB complex (pink) is superimposed with **A**) coprogen-bound FhuD from PDB: 1ESZ, **B**) albomycin-delta2-bound FhuD from PDB: 1K7S, and **C**) desferal-bound FhuD from PDB:1K2V. All the substrate-bound FhuD are colored in gray with the substrate shown as sticks.

**Figure S8.**
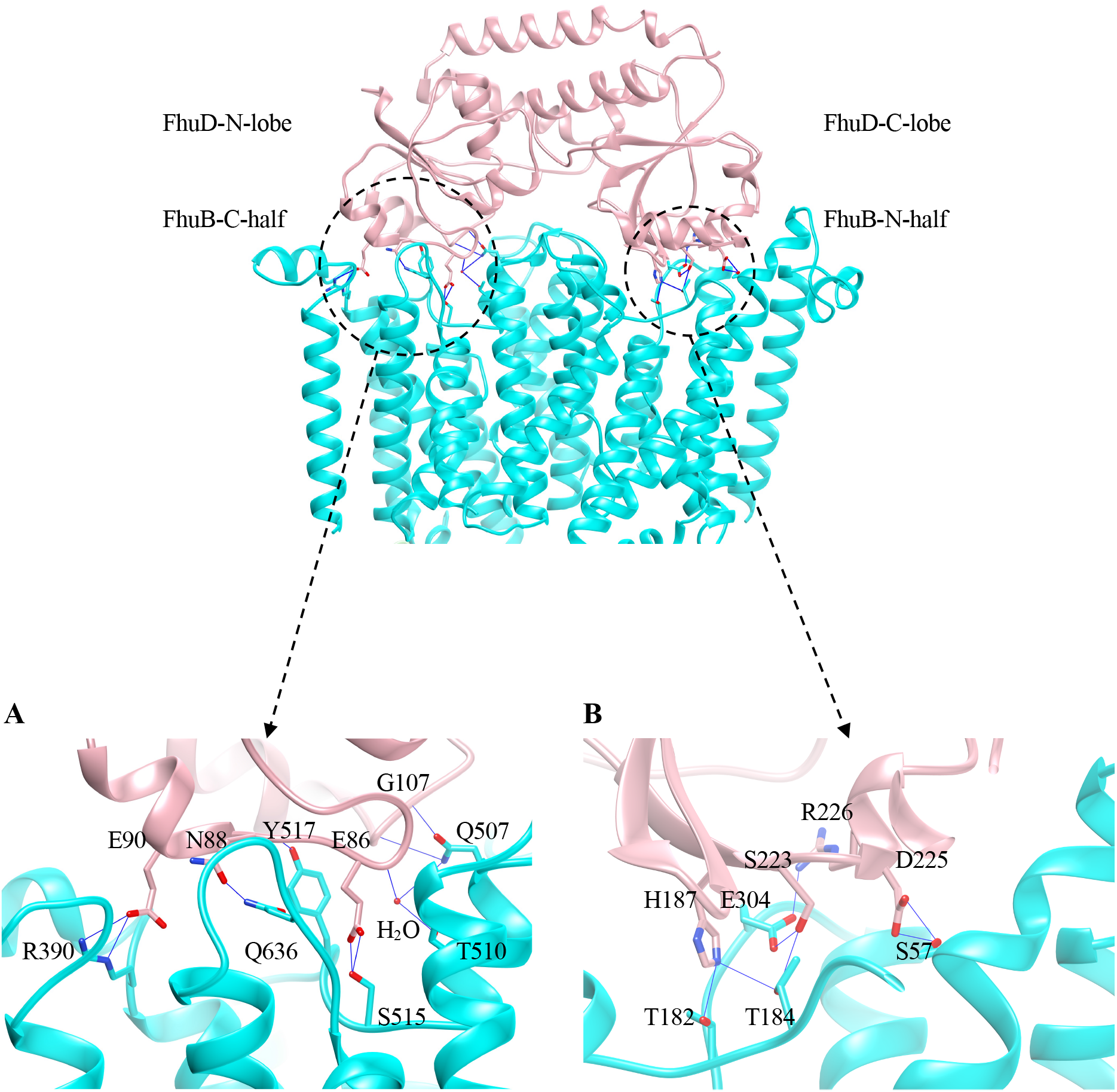
Interactions between FhuD and FhuB. **A**, Specific interactions between FhuD-C-lobe and FhuB-N-half. **B**, Specific interactions between FhuD-N-lobe and FhuB-C-half.

**Figure S9.**
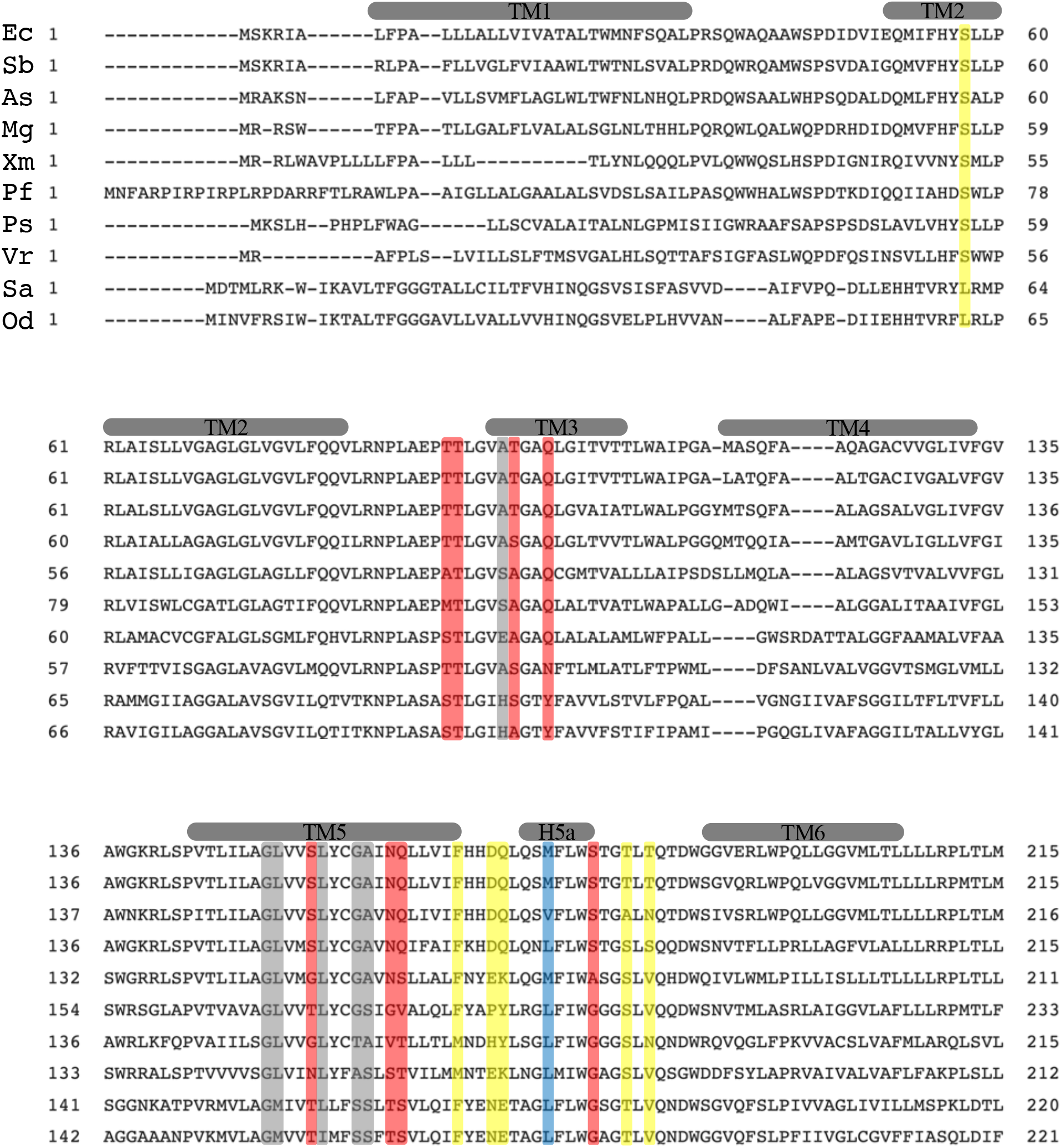
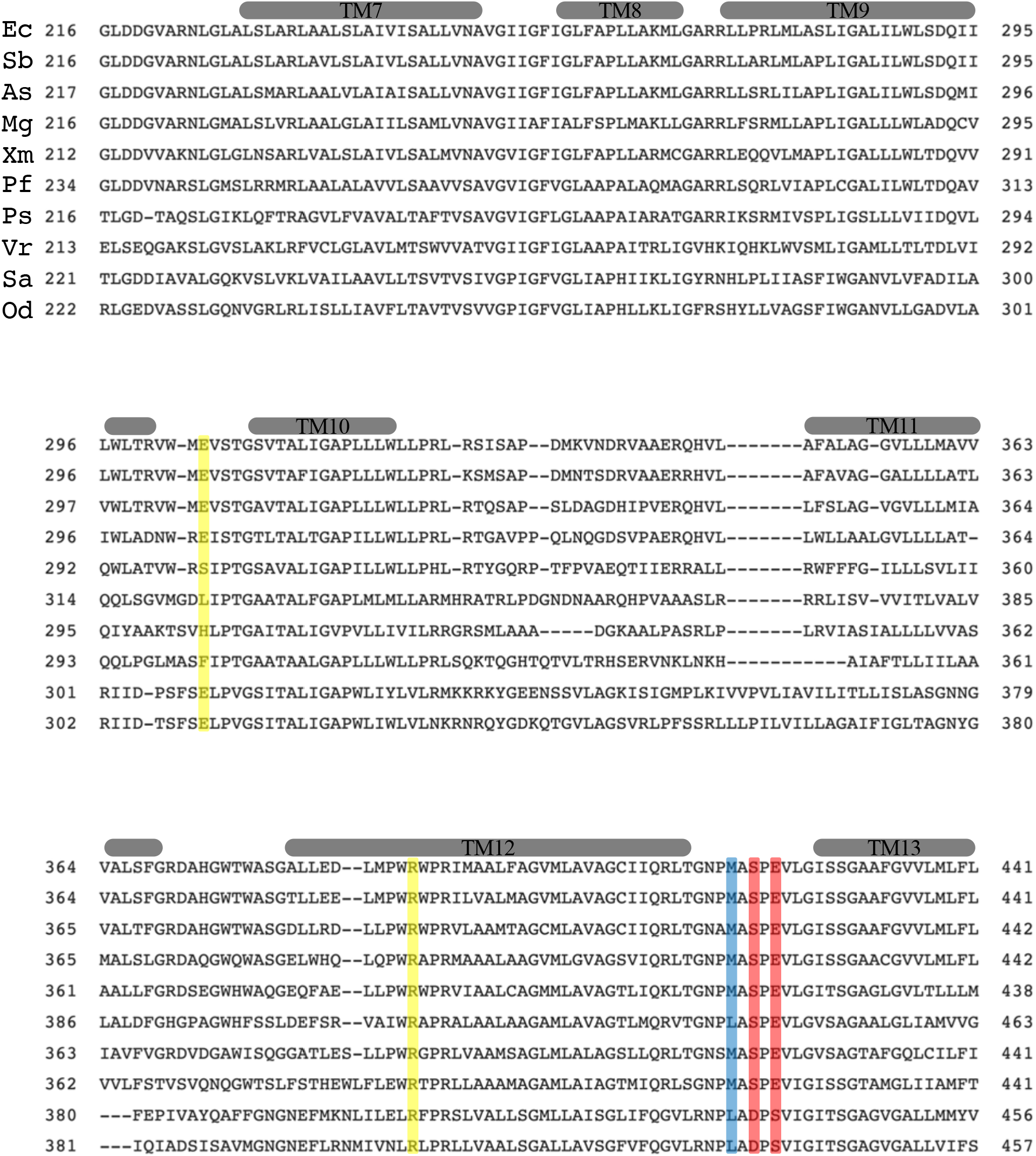
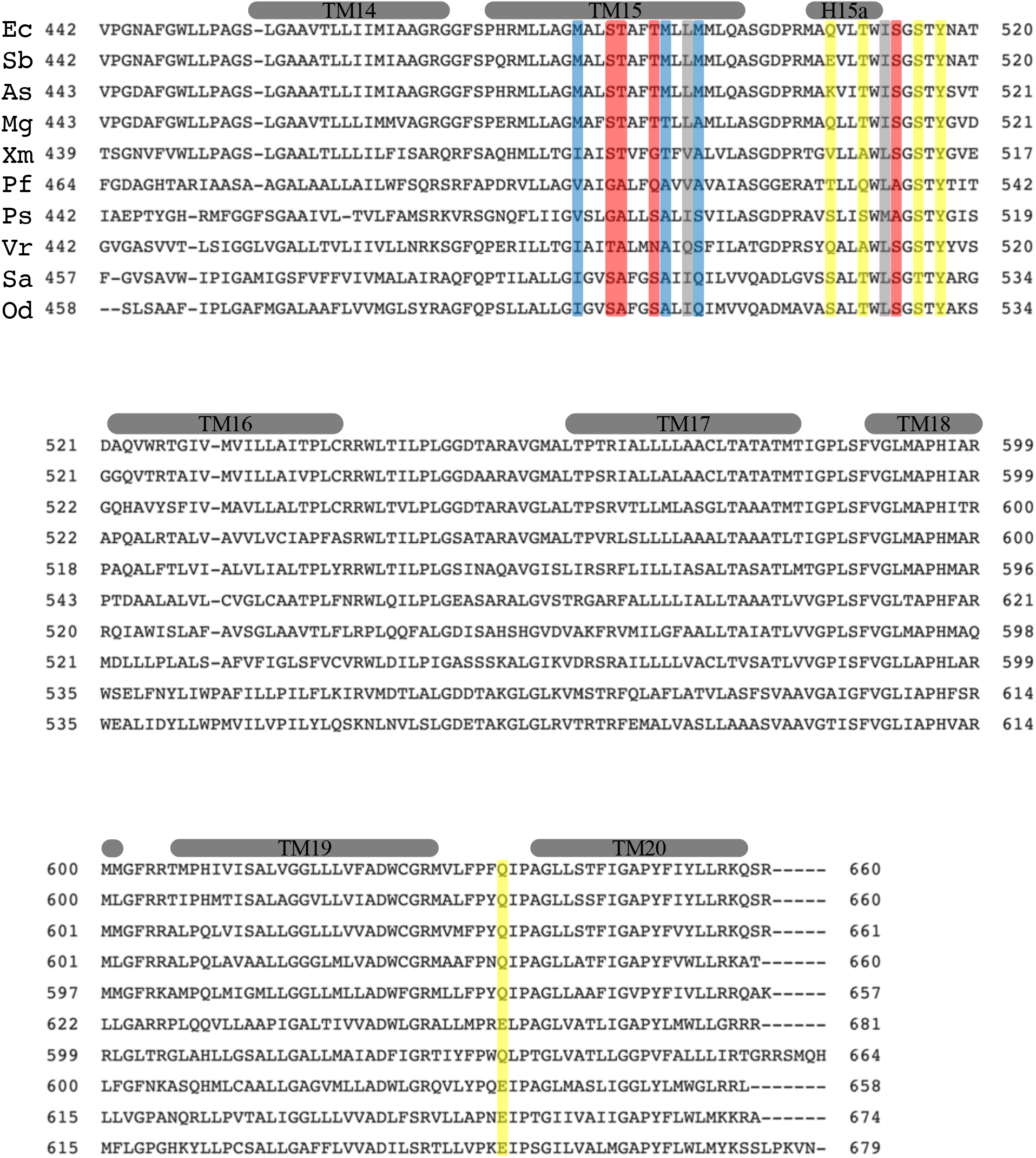
Multiple sequence alignment of FhuB from different species. The strains used and their sequence identity to *E. coli* FhuB are: *Salmonella bongori* (88.8%), *Atlantibacter subterranean* (80.5%), *Mixta gaviniae* (70.0%), *Xenorhabdus miraniensis* (60.2%), *Paraburkholderia fungorum* (49.7%), *Phyllobacterium sophorae (*40.7%), *Vibrio rotiferianus (*36.4%), *Sediminibacillus albus* (34.6%), *Oceanobacillus damuensis* (32.6%). Residues involved in FhuD-FhuB interaction are colored in yellow. Residues along the translocation pathway are colored in gray (small hydrophobic), red (polar) and blue (Mets).

**Figure S10.**
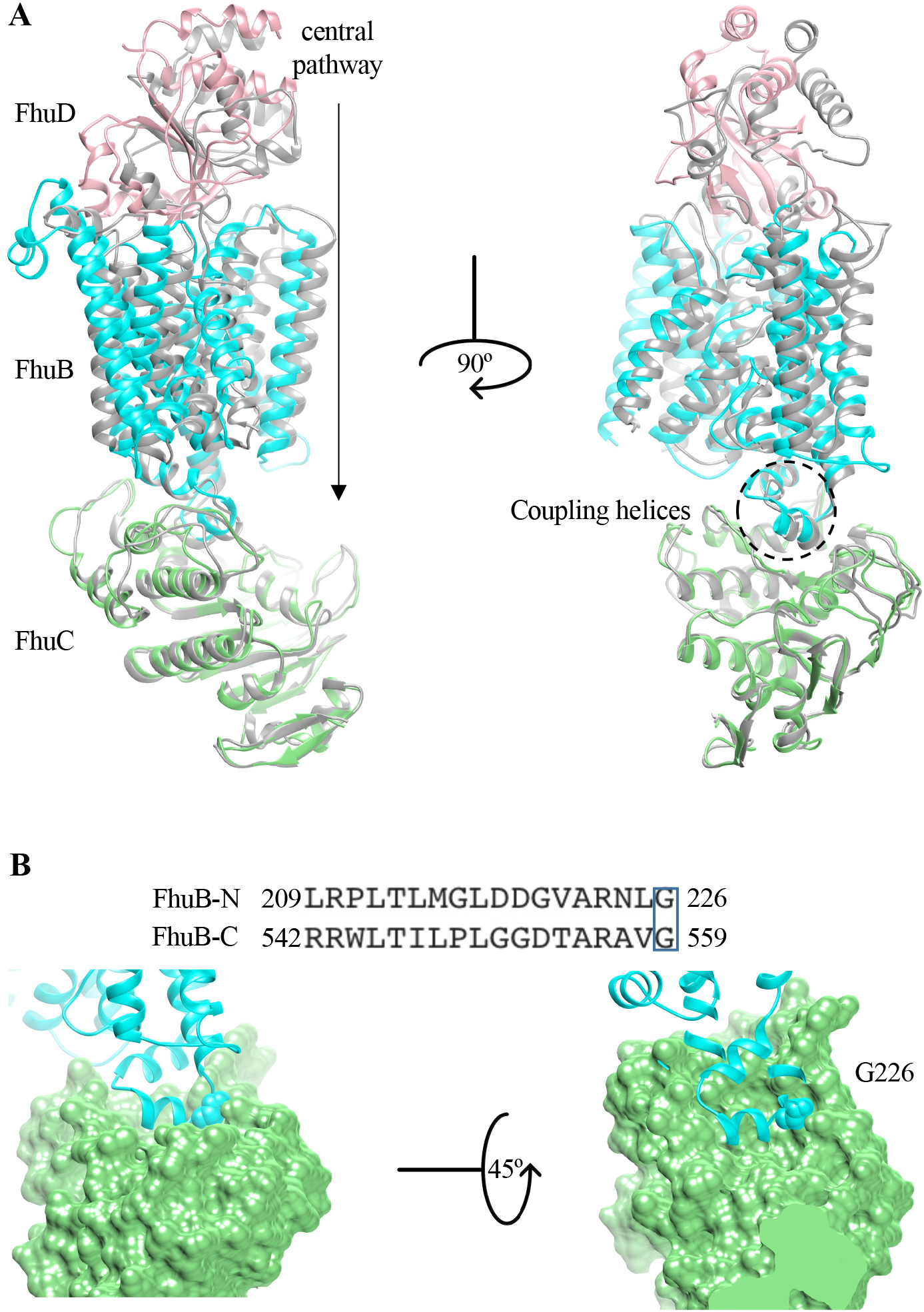
Association between FhuC and FhuB. **A**, Along the central channel in FhuB, the FhuCDB complex is divided into two halves and superimposed together with N-half multicolored and C-half in gray. **B,** Zoom in view of the FhuC-FhuB interface mediated by the coupling helices. Conserved residue FhuB-G226 (equivalent to FhuB-G559) is shown in ball mode, and the surface representation of FhuC is shown in green.

**Figure S11.**
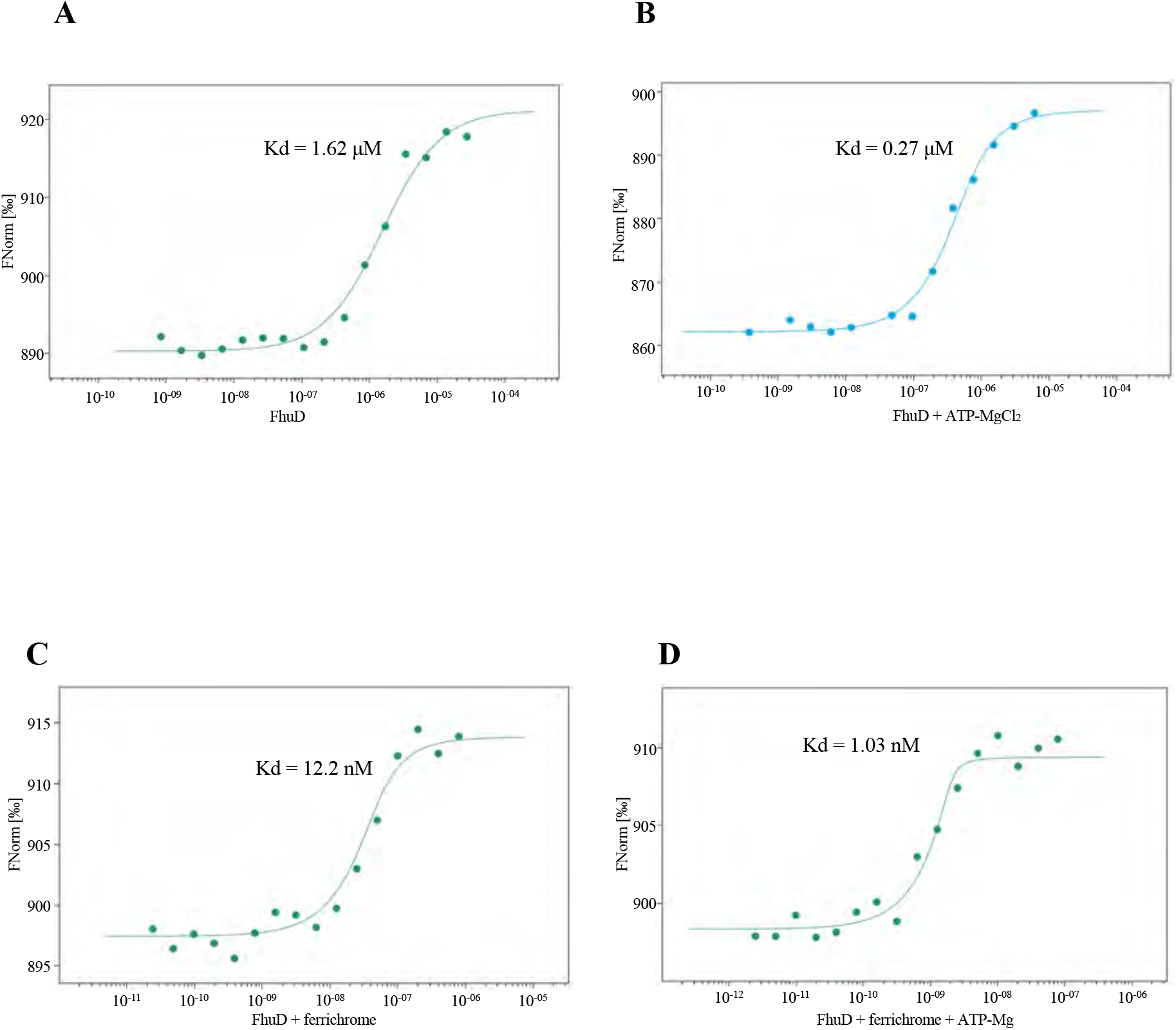
MST analysis of the binding between FhuCB and FhuD. **A**, FhuCB and FhuD. **B**, FhuCB and FhuD with ferrichrome. **C**, FhuCB and FhuD with ATP-MgCl_2_. **D**. FhuCB and FhuD with ferrichrome and ATP-MgCl_2_.

**Figure S12.**
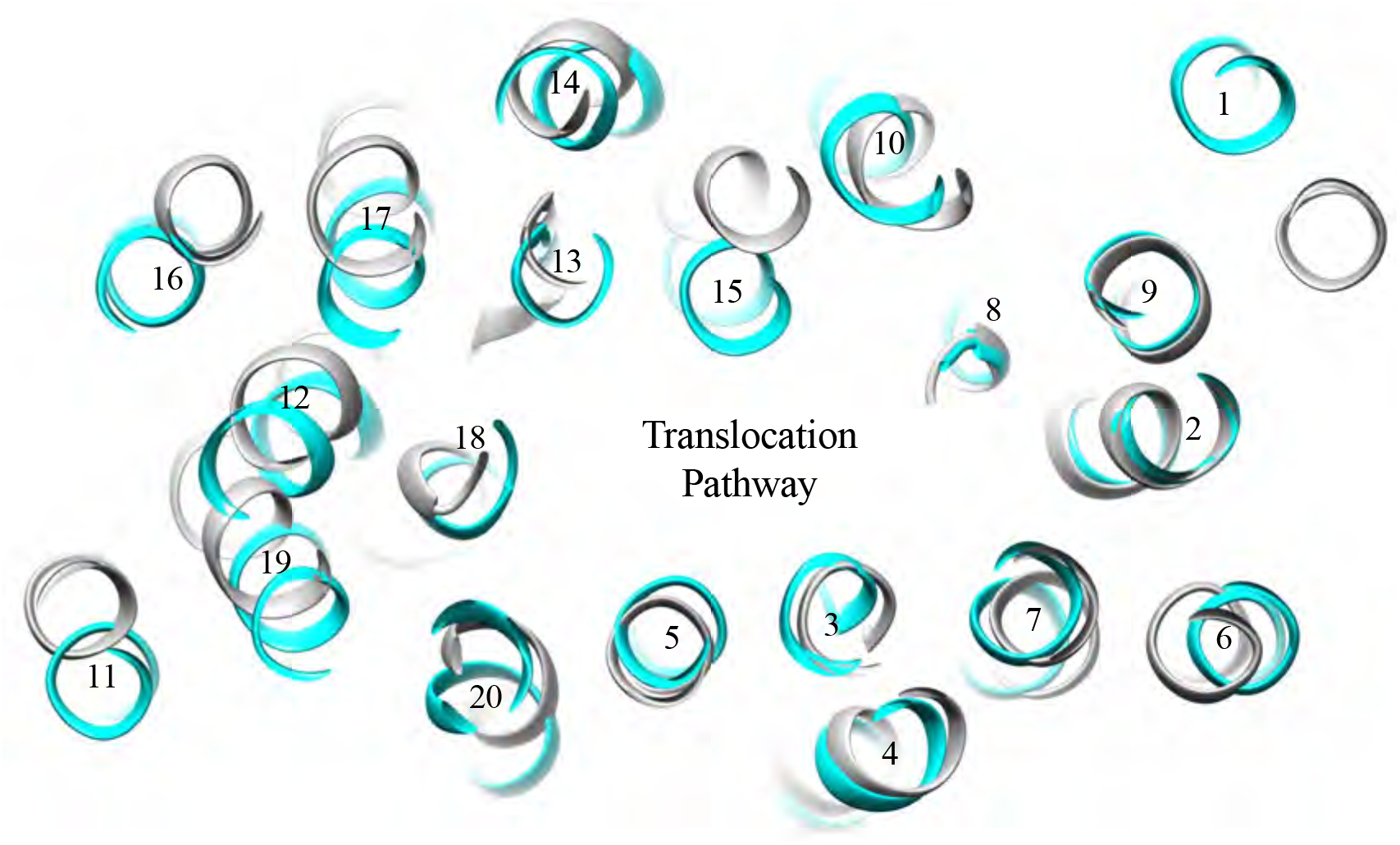
Comparison between FhuB and BhuU both in the inward-open conformation. FhuCDB is superimposed with BhuUVT (PDB:5B58). Looking down the translocation pathway from the periplasmic side, the sliced view of the TMs shows a narrower channel in FhuB (cyan) than BhuU (gray) because of the substantial movement of TM15.

**Figure S13.**
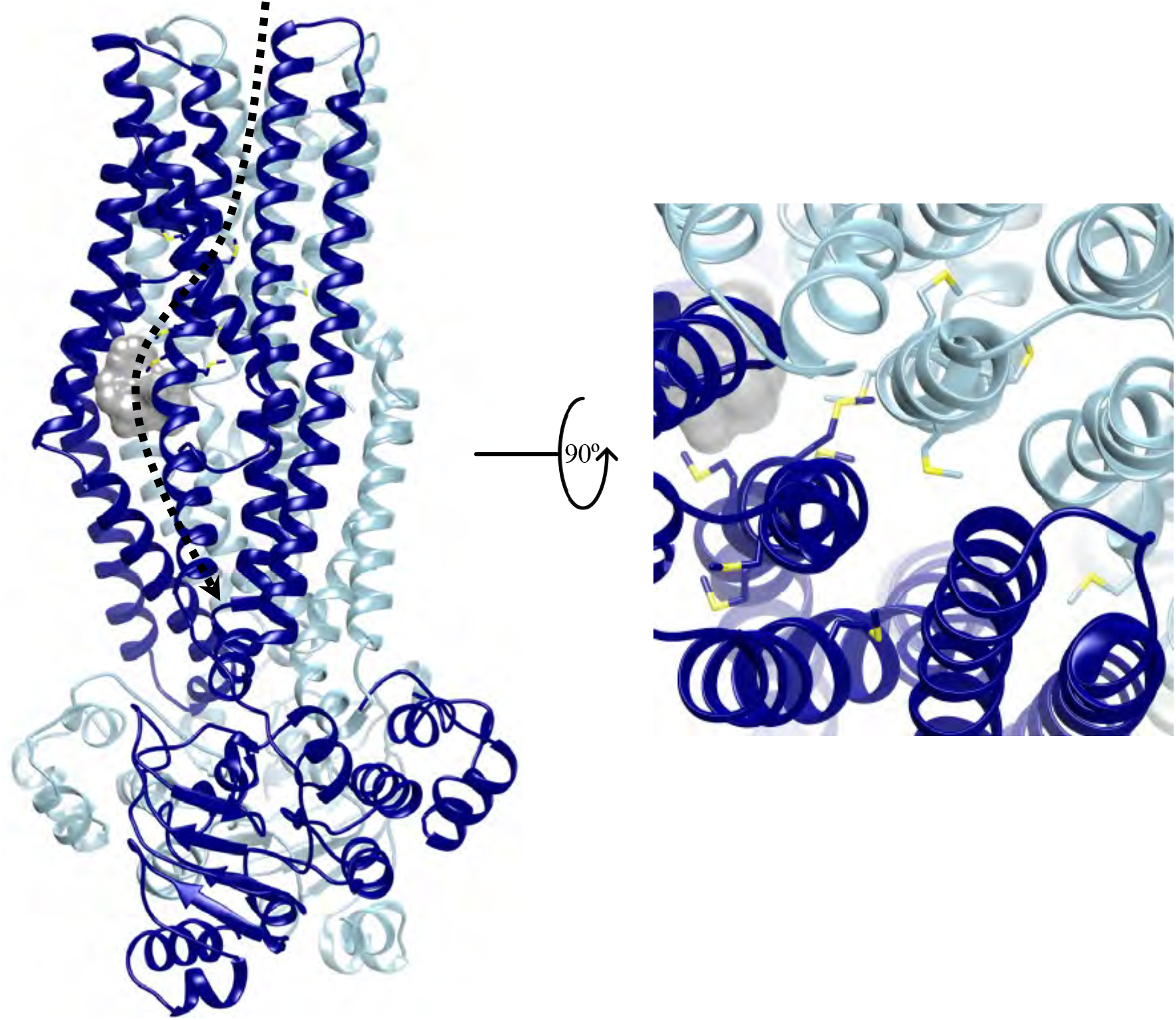
Methionine residues found in the structure of yersiniabactin importer YbtPQ (PDB:6P6J) around the central translocation pathway. YbtP is in dark blue, YbtQ is in light blue, the substrate Fe^3+^-yersiniabactin is in gray, Met residues are shown in stick mode and colored by heteroatom. The dotted line shows a putative translocation pathway.

**Figure S14.**
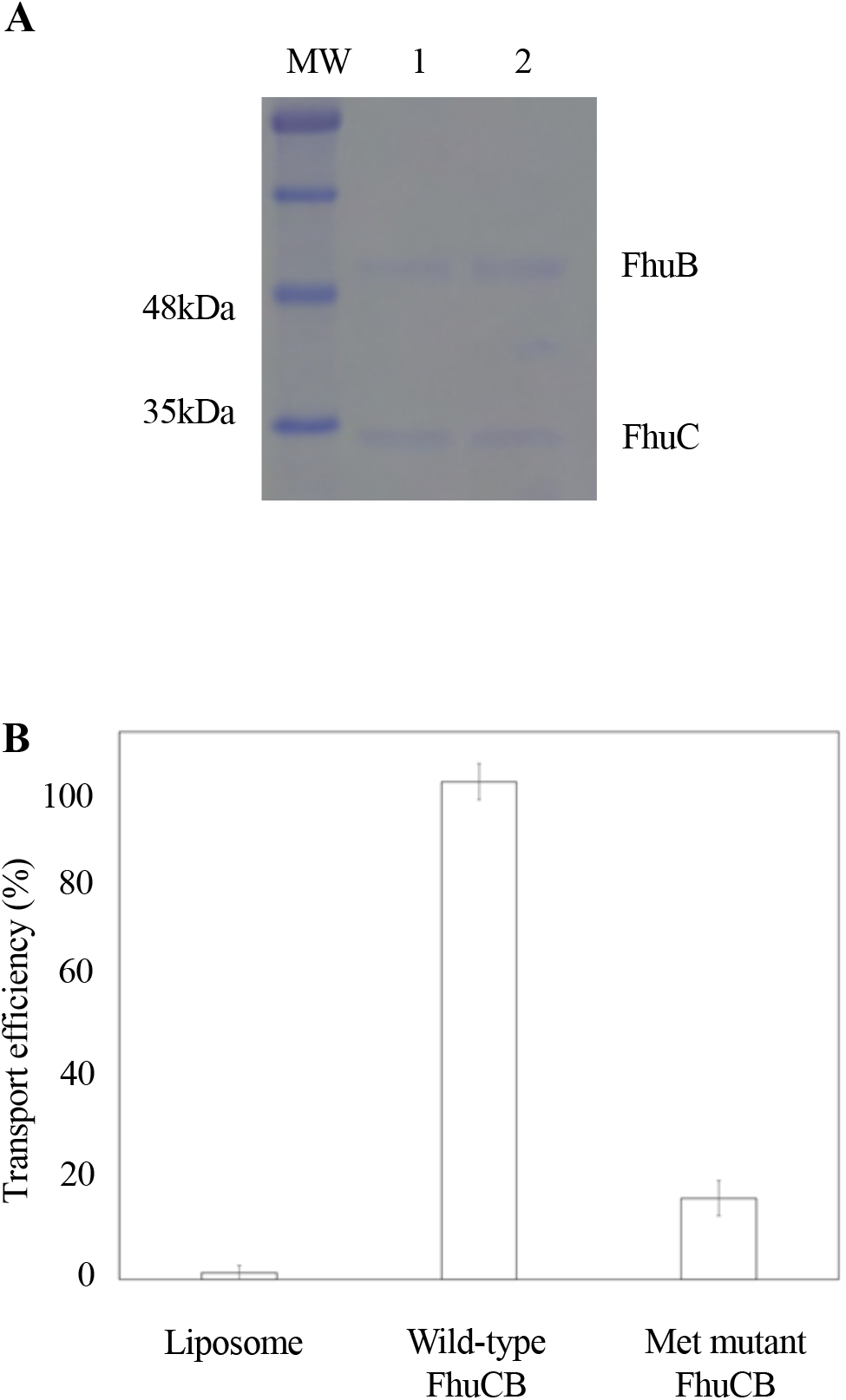
Substrate transport efficiency of FhuCB with Met mutations. **A,** Comparison of the amount of FhuCB reconstituted into liposomes. Lane 1: wild-type; lane 2: Met mutant. **B,** Liposomes without transporter show no transport, while proteoliposomes with Met mutations in FhuB show ~20% transport efficiency comparing to wild-type.

**Figure S15.**
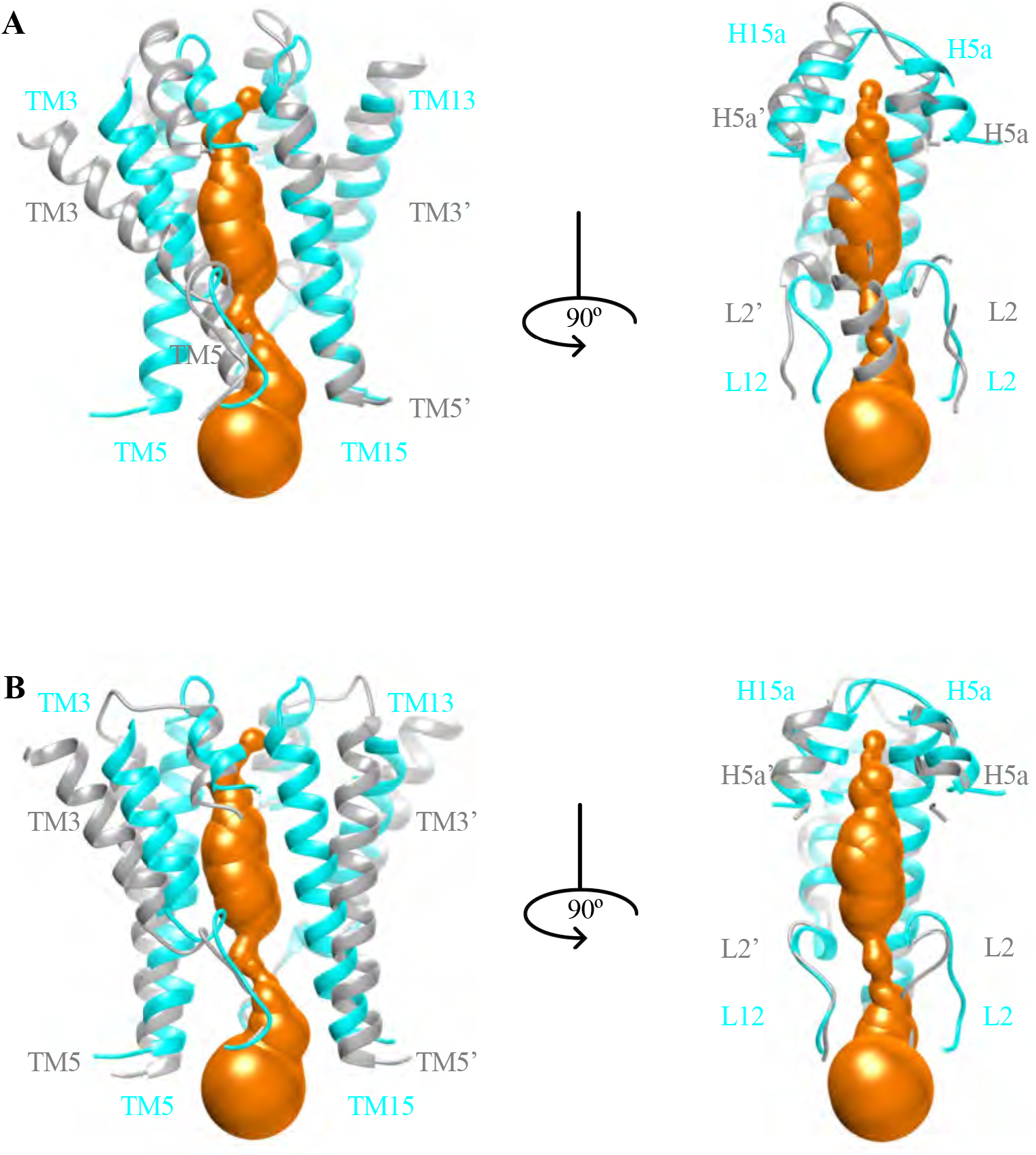
Comparison between FhuCDB and BtuCD structures. **A,** Closed cytoplasmic gate I (TM5 and its equivalent) in BtuCDF (2QI9, gray) is open in FhuB (cyan). **B**, Closed cytoplasmic gate II (L2 and its equivalent) in the outward-open BtuCD (4R9U, gray) is also open in FhuB (cyan).

**Table S1:**
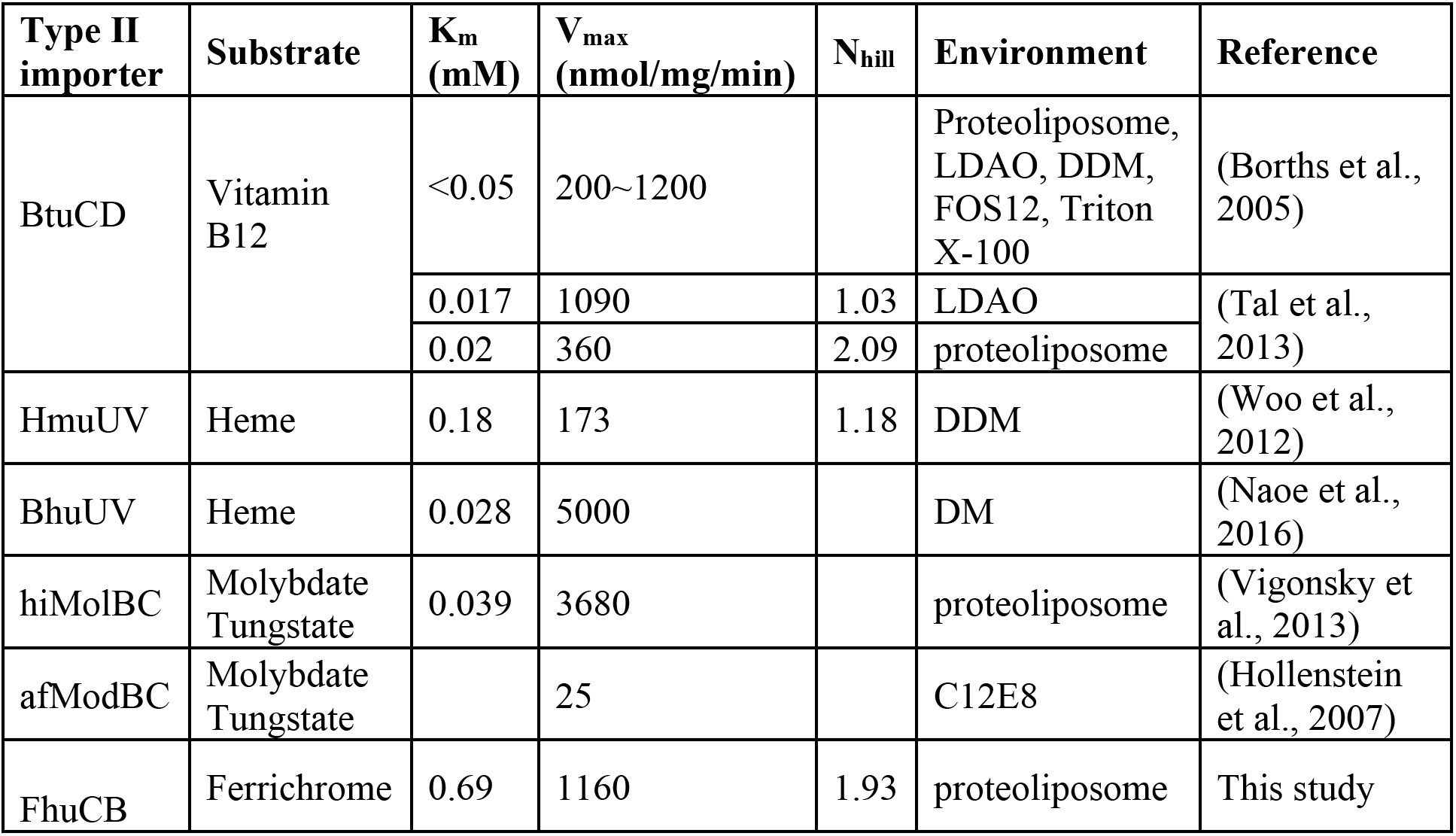
Kinetic constants of ATPase activity from known Type II importers.

**Table S2:** Cryo-EM data collection, refinement and validation statistics.

|  |  |
| --- | --- |
|  | FhuCDB in LMNG |
| **Data Collection and Processing** |  |
| Microscope | Titan Krios |
| Voltage (kV) | 300 |
| Magnification (nominal) | 92,000 |
| Electron Dose (e<sup>-</sup>/Å<sup>2</sup>) | 65 |
| Camera | Gatan K3 |
| Defocus range (um) | -1 ~ -2.5 |
| Pixel size (Å) | 0.399 (super resolution) |
| Movies collected | 9142 |
| Symmetry imposed | C1 |
| Final particle images (no.) | 128,131 |
| Map resolution (Å) | 3.4 |
| FSC cutoff | 0.143 |
| Map resolution range | 3.2 ~ 6.5 |
| Sharpening B-factor (Å<sup>2</sup>) | -106.7 |
| Software used to process data | cryoSPARC, RELION 3.1 |
| **Refinement statistics** |  |
| Initial model used | 1ESZ |
| Model resolution (FSC = 0.5) | 3.6 |
| Correlation Coefficient (Mask) | 0.76 |
| Number of protein atoms (non-H) | 10469 |
| Residues | 1405 |
| R.m.s. deviations |  |
| Bonds (Å) | 0.005 |
| Bond angles (°) | 0.715 |
| **Validation** |  |
| MolProbity score | 1.91 |
| Clash score | 10.40 |
| Poor rotamers (%) | 0.00 |
| **Ramachandran plot** |  |
| Favored (%) | 94.62 |
| Allowed (%) | 5.38 |
| Disallowed (%) | 0 |
| EMDB access code | EMD-23251 |
| PDB access code | 7LB8 |

**Table S3:** Interactions between FhuCB (wild-type and FhuB mutants) and FhuD analyzed using MST. All measured K_d_ values between FhuCB and FhuD are listed.

|  | FhuD | FhuD + ATP-Mg | FhuD-ferrichrome | FhuD-ferrichrome + ATP-Mg |
| --- | --- | --- | --- | --- |
| FhuCB wild-type | 1.62 μM | 0.27 μM | 12.2 nM | 1.03 nM |
| R390A |  |  |  | 75.8 nM |
| Q636A |  |  |  | 90.2 nM |
| Q507A,<br>T510A |  |  |  | 79.2 nM |
| S515A,<br>Y517A |  |  |  | Not detectable |
| D170A,<br>Q171A |  |  |  | Not detectable |
| S57A |  |  |  | 88.4 nM |
| E304A |  |  |  | 15.1 nM |
| T182A,<br>T184A |  |  |  | Not detectable |

